# Fire and forest loss in the Dominican Republic during the 21st Century

**DOI:** 10.1101/2021.06.15.448604

**Authors:** José-Ramón Martínez-Batlle

## Abstract

Forest loss is an environmental issue that threatens ecosystems in the Dominican Republic (the DR). Although shifting agriculture by slash-and-burn methods is thought to be the main driver of forest loss in the DR, empirical evidence of this relationship is still lacking. Since remotely sensed data on fire occurrence is a suitable proxy for estimating the spread of shifting agriculture, here I explore the association between forest loss and fire during the first 18 years of the 21st Century using zonal statistics and spatial autoregressive models on different spatio-temporal layouts. First, I found that both forest loss and fire were spatially autocorrelated and statistically associated with each other at a country scale over the study period, particularly in the western and central part of the DR. However, no statistical association between forest loss and fire was found in the eastern portion, a region that hosts a large international tourism hub. Second, deforestation and fire showed a joint cyclical variation pattern of approximately four years up to 2013, and from 2014 onwards deforestation alone followed a worrying upward trend, while at the same time fire activity declined significantly. Third, I found no significant differences in forest loss patterns between the deforested area of small (<1 ha) and large (>1 ha) clearings of forest. I propose these findings hold potential to inform land management policies that help reduce forest loss, particularly in protected areas, mountain areas, and the vicinity of tourism hubs.

## Introduction

Deforestation is a major concern for countries embracing the achievement of Sustainable Development Goal (SDG) 15 (Department of Economic and Social Affairs of the United Nations Secretariat, 2009; UN System Task Team on the Post-2015 UN Development Agenda, 2012). During the last decades, most countries have established reforestation programs to halt and reverse land degradation, but little effort has been made in preventing forest loss in preserved areas and secondary forests (Buřivalová et al., 2021; Heinrich et al., 2021). In addition, a conceptual framework for developing indicators for the SDG 15 is missing, making it hard to assess whether or not the goal is being met (Hák et al., 2016).

A global assessment of 21st-Century forest cover change, derived from Landsat satellite observations, was published in 2013 and has since been updated yearly (MC Hansen et al., 2013). Several research teams used the outcomes of Hansen et al.’s work to assess the changes and trends of forest cover in different countries and to explore the causes of deforestation (e.g., commodity-driven deforestation, shifting agriculture, and wildfires) (Curtis et al., 2018; Kalamandeen et al., 2018).

Despite the ecological importance of the forest ecosystems in the Dominican Republic (hereafter, the DR) (Cámara Artigas, 1997; Cano and Veloz, 2012; Hager and Zanoni, 1993; Olson et al., 2001), comprehensive assessments of forest loss are rare. The available evidence suggests that there is a close relationship between forest loss and shifting agriculture, the latter driven mainly by slash-and-burn practices (Cámara Artigas, 1997; Lloyd and León, 2019; Myers et al., 2004; OEA, 1967; Ovalle de Morel and Rodríguez Liriano, 1984; Tolentino and Peña, 1998; Wendell Werge, 1974; Zweifler et al., 1994). Although the Ministry of Agriculture and the National Bureau of Statistics of the DR have conducted agricultural censuses, their efforts have failed to provide consistent and spatially dense data on the intensity and extent of shifting agriculture activity over the last decades (ONE, 1982, 2016). Therefore, even a simple correlation analysis between forest loss and agricultural activity is unfeasible with the available data published by government institutions. A further limitation is the fact that traditional regression analysis cannot provide a systematic assessment of statistical association between variables that exhibit spatial autocorrelation, so spatial autoregressive models are needed (Anselin, 2013; RS Bivand et al., 2013).

Considering these limitations, I explore here the statistical associations between fire and forest loss in the DR in the first 18 years of the 21st Century, using spatial autoregressive models applied to public data remotely and consistently collected. Specifically, and referring to those 18 years, I answer the following questions: 1) Was fire statistically associated with forest loss and, if so, was fire a suitable predictor of forest loss? 2) Was there a greater degree of association of fire with small forest clearings than with larger ones? 3) Did spatial clustering or temporal trend of forest loss and fire exist? I hypothesize that both fire and forest loss were significantly and increasingly associated over time, that fire was a suitable predictor of forest loss regardless of the size of the clearings, and that both fire and forest loss were spatially autocorrelated over the study period.

This is the first study providing empirical evidence of the association between fire and forest loss in the DR. I assert that the results obtained increase knowledge on spatio-temporal patterns of forest loss. In addition, the findings could assist decision-makers in assessing the achievement of the SDGs, and in designing more effective policies for the long-term planning of nature conservation and for preventing wildfire and forest loss.

## Material and methods

### Data download and preparation

I used two types of datasets for this research (see Fig. 1): the collection of forest change layers from MC Hansen et al. (2013) and the fire point/hotspot locations from NASA (2019a,b). From the forest change data, I used the loss year and the tree cover thematic tiles, which I downloaded from the Global Forest Change 2000-2018 data service (Hansen/UMD/Google/USGS/NASA, 2019). The tree cover tiles classify the land area in tree canopy densities for the year 2000 as a baseline—where trees mean “vegetation taller than 5 m in height”— and the loss year tiles record the first year when the canopy reduced its density relative to the baseline year. Although Tropek et al. (2014) commented that the study underestimates forest loss, M Hansen et al. (2014) argued that such criticism is based on a misconception of the definition of forest used in their study.

**Figure 1.**
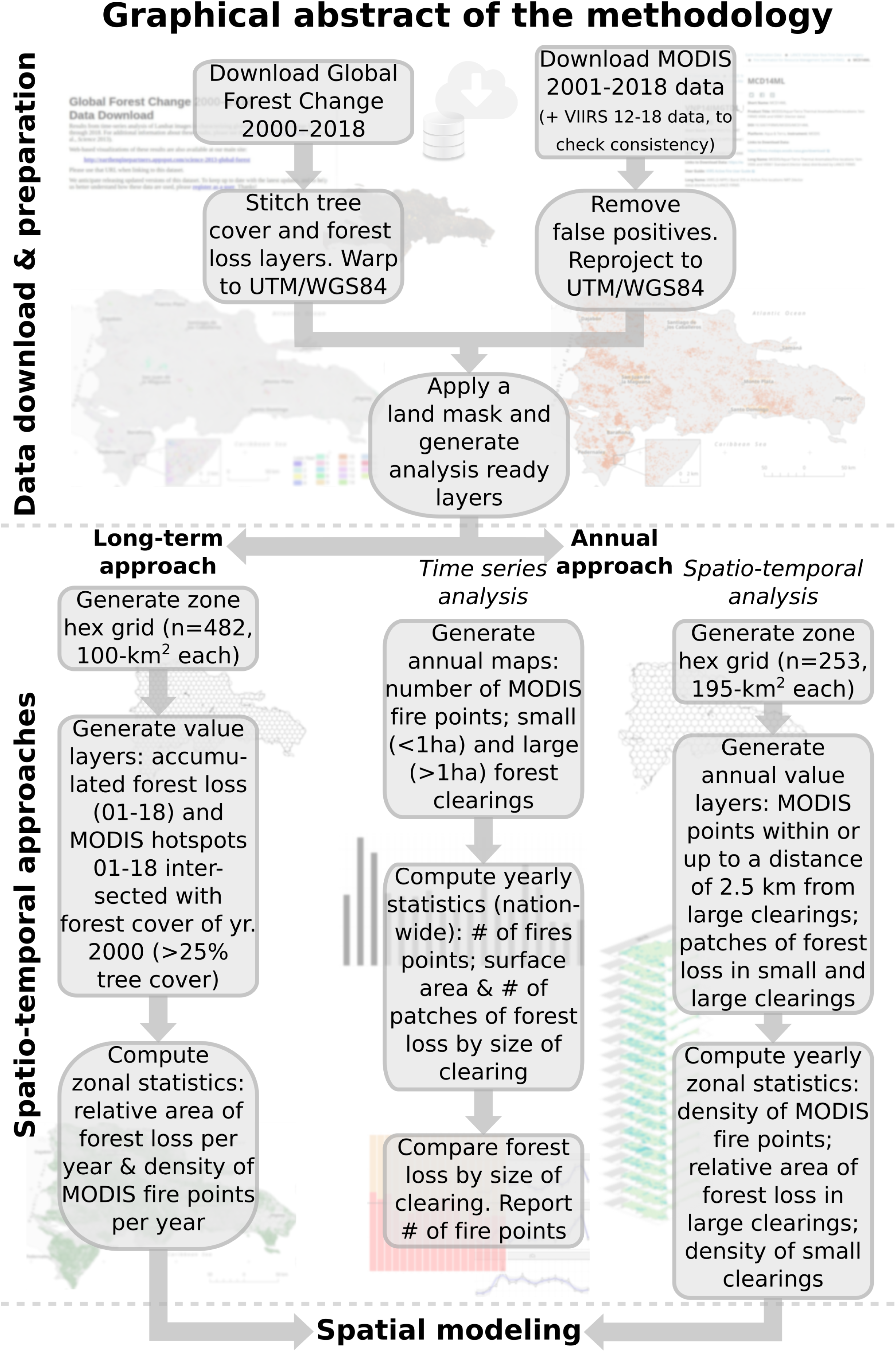
Graphical abstract of the methodology. See text for details and see the Data and code availability section for provided scripts.

I stitched together the tiles from these datasets to form a seamless mosaic, and then warped the results on to the UTM/WGS84 datum, from which I later produced continuous maps of the DR mainland territory by masking out the ocean/lake areas (Fig. 2). Since these products do not distinguish plantations (e.g., oil palm and avocado plantations) from forest, I acknowledged this limitation when running exploratory analysis and building spatial models.

**Figure 2.**
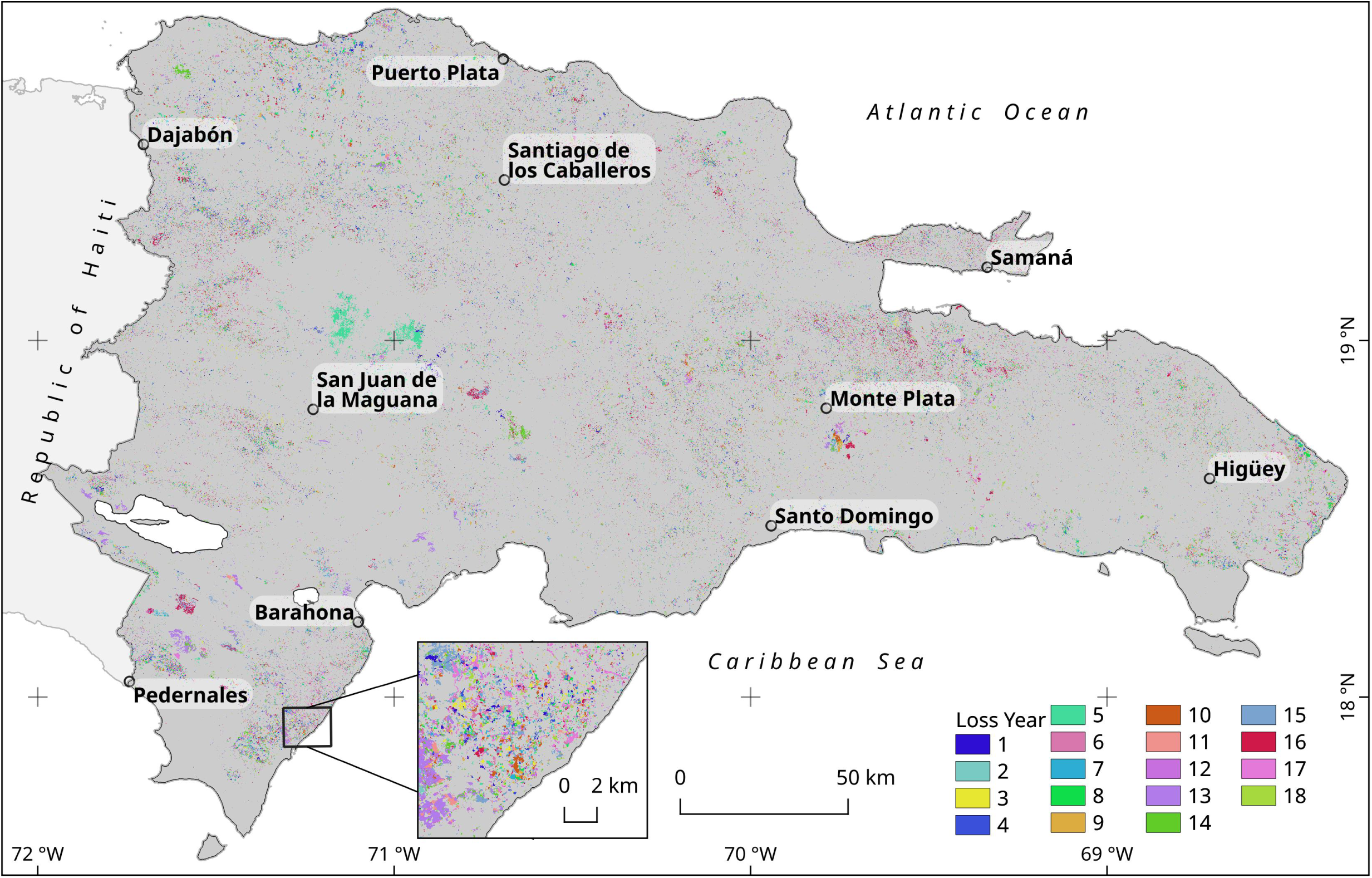
Loss year layer from 2001 to 2018 for the Dominican Republic, according to MC Hansen et al. (2013). Labelled points denote the location of some cities chosen as reference.

Moreover, the fire/hotspot data consisted of two products of the NASA’s Fire Information for Resource Management System (FIRMS) processed by the University of Maryland, provided as point layer files by the LANCE/ESDIS platform, covering two overlapping periods of time (NASA, 2019a,b). The most comprehensive dataset, labeled as “MODIS Collection 6 standard quality Thermal Anomalies / Fire locations” (MCD14ML), comprised fire data from 2000 to 2018. The second product, labeled as “VIIRS 375 m standard Active Fire and Thermal Anomalies product” (VNP14IMGTML), comprised locations of fires and thermal anomalies since 2012 up to the present time.

Since the MODIS dataset covered the longest time period, I used it as the reference database for the multiyear analyses. Furthermore, considering that the VIIRS time series described just the last third of the analyzed period, I used this set for assessing the consistency and sensitivity of the MODIS data. To do this, I generated a subset of the MODIS and VIIRS datasets from the 2012-2018 period, summarizing the number of fire points per month. With this subset, I performed a cross-correlation analysis and fitted a linear model using the number of MODIS fire points per month as the independent variable and the number of VIIRS fire points as the response variable (Venables and Ripley, 2002). After the consistency check, I used only the MODIS dataset for further analyses.

The FIRMS source web service states that there are missing data at known dates in the MODIS product, but, since this issue affects a minimal portion of the time series—few days of 2001, 2002, 2007 and 2009, overall, less than 30 days—, I decided to acknowledge it and use the entire dataset without applying missing data algorithms.

Most of the fire data points from the FIRMS collections accounted for actual fires and thermal anomalies, but there were also noisy records (e.g., false positives) that could affect the results. Thus, I removed the persistent thermal anomalies records with little or no potential to produce wildfires, such as those originating from landfills with spontaneous combustion and industrial furnaces. I refer to the resulting outcome as “the noise-free versions of the fire points datasets” or simply “the noise-free versions” (see Supplementary Information section and Fig. S1 for details). For consistency reasons, I reprojected the point data files to conform to the UTM/WGS84 datum. Last, I applied a mask comprising the DR land area to each dataset used in the study.

### Spatio-temporal approaches

I used two different spatio-temporal approaches to answer the questions posed in this study, which I refer to as “the long-term approach” and “the annual approach”, respectively (Fig. 1). In both approaches, I applied spatial statistical techniques to explore association patterns between forest loss and fire, using statistical summaries generated from zonal grids and value layers.

#### Long-term approach

In this approach, I assessed the association between forest loss and fire in the study period—2001-2018— using a zonal grid. I focused the analysis on the areas with 25% or higher tree cover in year 2000 as a baseline, which I refer to as “forest cover in 2000”, or simply “forest cover” (Fig. 3). I used this baseline for two reasons: 1) The 2000 tree cover serves as a baseline for global forest change studies (MC Hansen et al., 2013); 2) 25% tree cover is an appropriate threshold to cover different vegetation types, including tropical semi-deciduous and seasonally dry forests.

**Figure 3.**
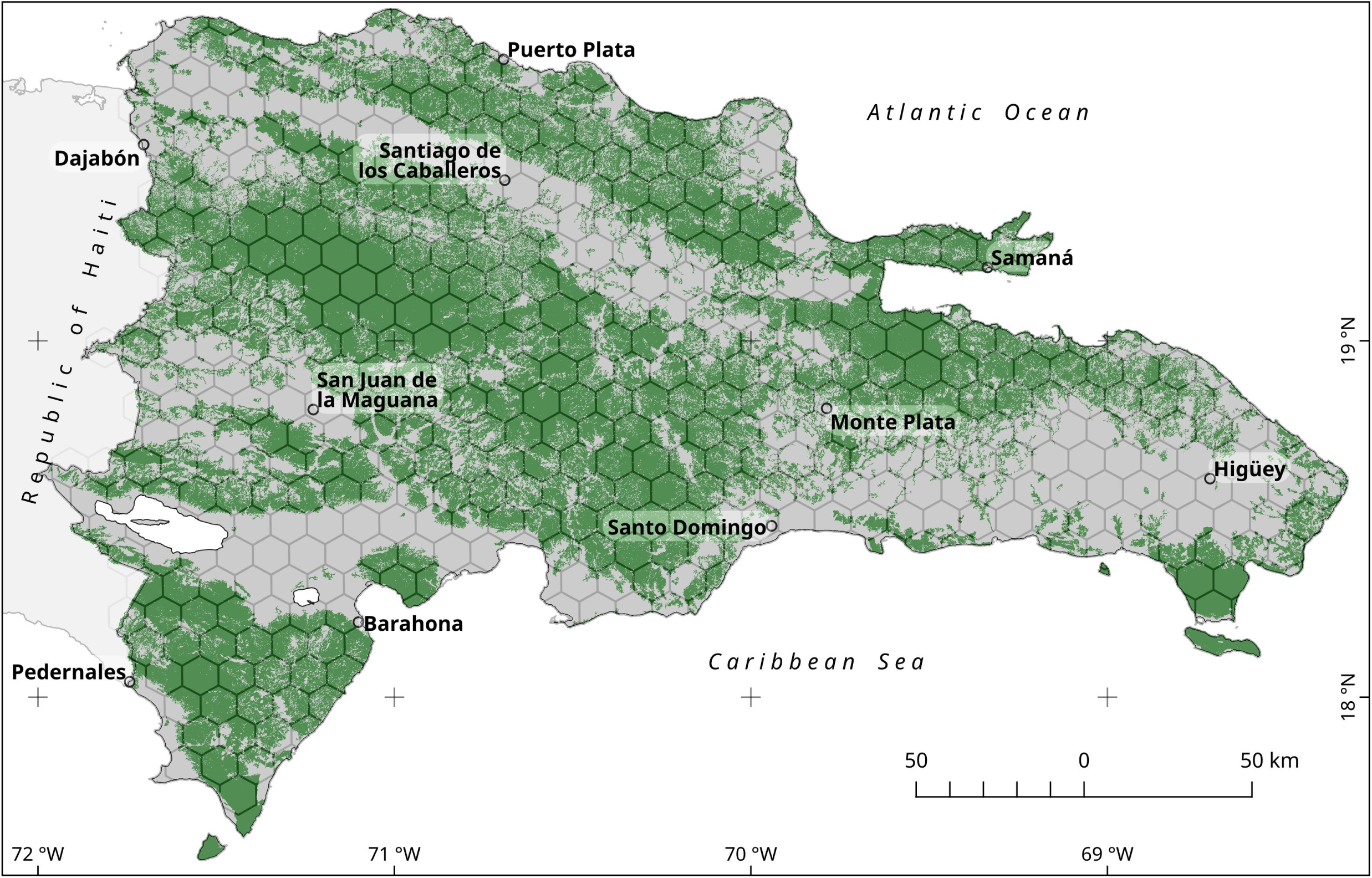
DR forest cover in the year 2000. Areas with a canopy closure equal to or greater than 25% in tree cover map of MC Hansen et al. (2013) were classified as forest. The hexagonal grid overlaid was used for zonal statistics computations of the long-term approach. See text for details.

The zonal grid created for this approach consisted of 482 adjacent hexagons, each with a nominal surface area of 100 km^2^ and having at least 45% of its area on mainland territory (Fig. 3). With this setting, the total area of the zonal grid was approximately 46,200 km^2^, which is indeed slightly smaller than the DR territory (approximately 48,400 km^2^).

To generate the fire data layer, I used the noise-free version of the MODIS dataset as input and selected the fire points falling into the above defined forest cover. Then, I computed the number of fire points for each hexagon of the zonal grid. Last, I divided the number of points in each cell by its area (in km^2^) and by the number of years, which resulted in fire density (Fig. 1).

Moreover, to generate the forest loss data layer, I pooled forest loss surface area representing the period 2001 to 2018, then divided it by the corresponding cell size and by 18 years to obtain the average forest loss per unit area per year

While the long-term approach provides a useful summary of the relationships between fire density and forest loss for the period analyzed, most of the trends and other insightful patterns would remain unknown without an annual analytical approach.

#### Annual approach

For this approach, I analyzed temporal trends and statistical association between forest loss and fire on an annual basis with time series and spatio-temporal analyses.

I used the forest loss year raster to generate 18 annual maps. From each map, I grouped the connected cells belonging to the same patch using the Queen’s case neighborhood, and then calculated the surface area of the clumped patches. Additionally, I produced annual maps of “small forest clearings” with patches of less than 1 ha in size, and maps of “medium- and large-sized forest clearings” (or “large clearings”) with patches larger than 1 ha (Fig. 1). Then, I computed the annual forest loss separately by size of clearing, summing up the surface area values of the individual patches of each loss map, and assessed the homogeneity of annual average values using a paired *t*-test. Finally, I used the annual data to generate a time series of forest loss and fire occurrence, from which I extracted the trend and cyclical components using the “Christiano-Fitzgerald” and “Hodrick-Prescott” filters (Balcilar, 2019).

To perform the spatio-temporal analysis, I summarized the annual forest loss and fire density over a regular hexagon grid of 253 hexagons, each of which had a maximum area of approximately 195 km^2^. This larger area than for cells used in the long-term approach was chosen to reduce the skewness of variables distributions and improve adherence to normality. Afterward, I performed a zonal statistical analysis of forest loss, using separate metrics for large and small clearings. For large clearings, I used the relative area of annual forest loss (measured in km^2^ per 100 km^2^), since that metric is suitable for characterizing the deforestation activity on a given cell. For small clearings, density of patches (measured in number of patches per 100 km^2^) was used, since the relative area may be irrelevant for summarizing small clearings on a given cell.

Finally, to obtain the yearly subsets of fire points, I used the noise-free version of the MODIS dataset for the 2001-2018 period. I generated annual maps of fire points using the date field of the dataset. Then, from the annual maps of large clearings, buffer zones were created around the patches at a maximum distance of 2.5 km. Afterward, I generated the corresponding annual subsets of fire points, selecting only those falling within the patches and/or their buffer zones (Fig. S6). Last, I summarized, over the hexagon grid, the yearly density of MODIS fire points per 100 km^2^.

### Spatial modeling

For both the long-term and annual approaches, I conducted exploratory spatial data analysis (ESDA) and fitted several models using maximum likelihood estimation. First, I assessed the normality of the variables using Shapiro-Wilk tests and QQ plots, and applied Tukey’s Ladder of Power transformations to those variables departing from normality before performing spatial analysis, in order to fulfill the normality assumption or to reduce the skewness of the variables (Mangiafico, 2019).

For each of the grids used in this study, I created neighbour objects between hexagons based on the criterion of contiguity. As expected, each hexagon became the neighbour of six other contiguous hexagons, except for those located at the edge of the grid. Then, I defined spatial weights from the neighbour objects using the “W-style”—row standardization—, in which the weights of all the neighbour relationships for each areal unit summed 1.

As a prerequisite for modeling, I tested whether fire density and forest loss variables showed spatial autocorrelation, using Moran scatterplots and Moran’s *I* tests (Sokal and Oden, 1978). I also generated local indicators of spatial association maps (hereafter “LISA maps”), to represent high-high and low-low clusters of fire density and forest loss across the DR (Anselin, 1995, 1996; Anselin and Rey, 2010; R Bivand, Altman, et al., 2017; R Bivand, Hauke, et al., 2013; R Bivand and Piras, 2015; R Bivand and Wong, 2018; RS Bivand et al., 2013). A high-high cluster—hereafter HH cluster—is a group of cells in which high values are surrounded primarily by other high values. Conversely, a low-low cluster—hereafter LL cluster—is a group of cells with low values surrounded by other low values.

Since both fire density and forest loss variables showed significant patterns of spatial autocorrelation, I analyzed the statistical association between them using spatial autoregressive models. Specifically, I fitted spatial lag and spatial error models using fire density as a predictor variable and forest loss as a response variable. In the long-term approach, I evaluated the prediction performance of spatial lag and spatial error models. The most suitable model was chosen based on the results of the Lagrange Multiplier diagnostic for spatial dependence in linear models, the results of the Breusch-Pagan test for heteroskedasticity of residuals, and the Akaike information criterion (AIC) (Anselin, 2013; RS Bivand et al., 2013; Breusch and Pagan, 1979; LeSage, 2015; Sakamoto et al., 1986). In the annual approach, I generated yearly spatial error models to assess the statistical association between fire and forest loss. In general, and unless otherwise indicated, for all statistical tests, I used a significance level *α* = 0.05, and for error estimation I used a 95% confidence level.

All the results, including statistical summaries, maps, and graphics were produced with QGIS and R programming environment, using parallel computing packages for generating the zonal statistics outcomes, as well as multiple packages for data visualization and spatial modeling (Greenberg and Mattiuzzi, 2018; Hijmans, 2019; Kuhn et al., 2019; Pebesma, 2018, 2019; QGIS Development Team, 2020; R Core Team, 2020; Tennekes, 2018; Venables and Ripley, 2002; Weston, 2019; Wickham, 2017) (see also the Data, script and code availability section).

## Results

### MODIS data consistency and sensitivity

The MODIS dataset showed high consistency with the VIIRS dataset for the period 2012-2018. The cross-correlation between the two time series was close to 1 and positive for lag 0 (see Supplementary Information section for details). In terms of sensitivity, for every fire point detected by the MODIS sensor on a monthly basis, the VIIRS sensor detected six times more hotspots. Likewise, the latter detected an average of eight points that were not detected by the former. However, although the sensitivity of the MODIS sensor was lower than that of the VIIRS sensor—which was expected given its lower resolution—, its performance was highly correlated across time and thus appropriate for the purposes of this study.

### Long-term analytical approach of forest loss and fire

#### Overall statistics

The surface area of forest loss relative to the forest cover in the year 2000, was approximately 3,100 km^2^ during the period 2001-2018, which represents c. 7% of the entire grid analyzed (Table 1). Moreover, during the same period, the MODIS sensor recorded almost 11,700 points within forest cover areas.

**Table 1.** Forest loss and number of fire points within forest cover, summarized using a grid of 482 hexagons, for the period 2001-2018. The baseline year for the forest is 2000^†^.

| Attribute | Period 2001-2018<br>(fire data from MODIS) |
| --- | --- |
| Total number of fire points | 11,666 |
| Average number of fire points per 100 km<sup>2</sup> | 25.13 |
| Average number of fire points per 100 km<sup>2</sup> per year | 1.4 |
| Maximum number of fire points per 100 km<sup>2</sup> per year | 13.22 |
| Total forest loss area in km<sup>2</sup> (approximate percentage relative to the entire grid) | 3135.22 (6.8%) |
| Average forest loss area (km<sup>2</sup>) per 100 km<sup>2</sup> | 6.72 |
| Average forest loss area (km<sup>2</sup>) per 100 km<sup>2</sup> per year | 0.37 |
| Maximum forest loss area (km<sup>2</sup>) per 100 km<sup>2</sup> per year | 1.82 |

<sup>†</sup> The values of this table were summarized using zonal statistics techniques relative to a hexagonal grid. Thus, actual values of the entire DR may be slightly larger, since forest loss patches and fire points outside the grid were ignored.

#### Spatial patterns

Most of the DR mainland territory experienced low levels of forest loss from 2001 to 2018 (i.e., < 6 km^2^ per 100 km^2^). However, high levels of forest loss were common in several mountain ranges and protected areas, such as Los Haitises karst region, Samaná Peninsula, Sierra de Bahoruco, and the Cordillera Central southern and northwestern borders (Fig. 4-A). It should be particularly emphasized that inaccessible areas in Los Haitises, Sierra de Bahoruco and southern Cordillera Central, reached worrisome records of forest loss greater than 25 km^2^ per 100 km^2^. Additionally, the Eastern Region—Punta Cana and its surroundings, where tourism development has grown steadily since the 1990s—experienced high rates of forest loss during this period.

**Figure 4.**
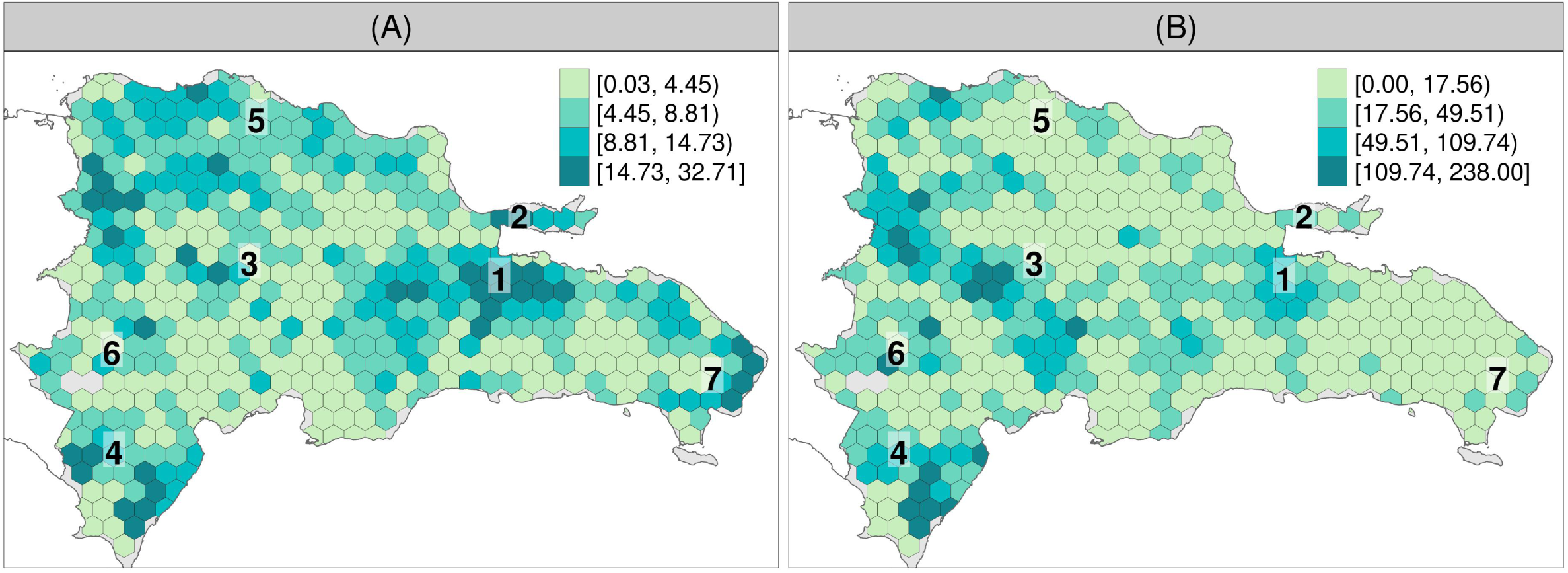
(A) Forest loss (in km^2^ per 100 km^2^) for the period 2001-2018. (B) Number of fire points per 100 km^2^ within forest cover for the period 2001-2018 using MODIS dataset. The baseline year for forest cover is 2000. Reference locations: 1 Los Haitises; 2 Samaná Peninsula; 3 Cordillera Central mountain range; 4 Sierra de Bahoruco; 5 Cordillera Septentrional; 6 Sierra de Neyba; 7 Eastern Region.

Furthermore, the density of fire points in the 2001-2018 period showed a distribution pattern similar to that of forest loss. High densities of fire points were fairly common in many areas, such as the southern margin of Cordillera Central, Sierra de Bahoruco, Sierra de Neyba, and Los Haitises, with more than 30 fire points per 100 km^2^ detected by the MODIS sensor (Fig. 4-B).

The analyses of the spatial autocorrelation of the transformed variables consistently showed the presence of positive autocorrelation patterns (Table S1, Figs. 5 and S5). The prevalence of HH clusters indicates that forest loss was notably widespread during the period 2001-2018 in Los Haitises, Sierra de Bahoruco, Samaná Peninsula, and the Eastern Region (Fig. 5-A).

**Figure 5.**
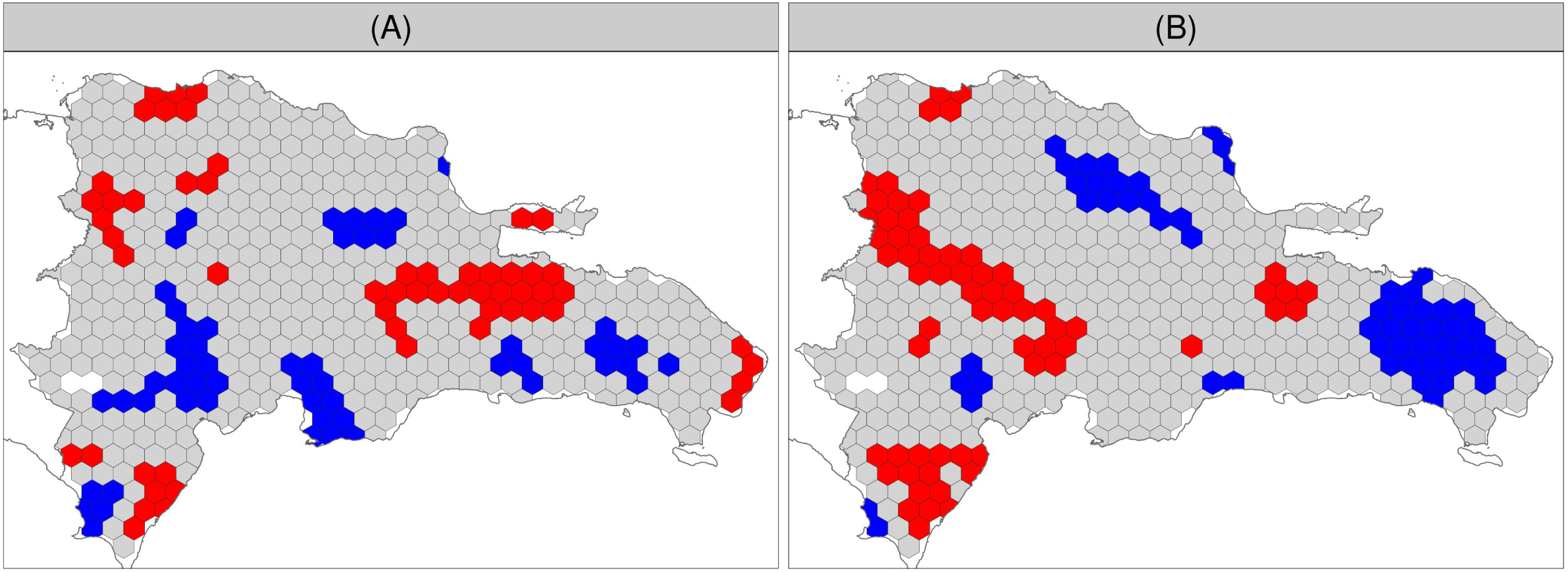
LISA maps (local indicators of spatial association maps) of (A) forest loss per unit area averaged per year for the period 2001-2018, and (B) fire points per km^2^ averaged per year within forest cover for the same periods using the MODIS dataset. The Tukey’s Ladder of Powers transformed versions of the variables were used as inputs in all cases. Each hexagon was classified either as HH cluster—group of cells in which high values are surrounded by other high values—(depicted in red), LL cluster—cells with low values surrounded by other low values—(depicted in blue), or no significant spatial association (grey) regarding the corresponding variable.

Moreover, HH clusters of fire density were notably widespread in the southern margin of Cordillera Central, Los Haitises, western Cordillera Septentrional, Sierra de Neyba, and Sierra de Bahoruco (Fig. 5-B). There is a noticeable high degree of agreement between forest loss and fire density LISA maps in parts of Los Haitises, Sierra de Bahoruco, and other areas, suggesting that an association exists between these variables. A notable exception is the Eastern Region, where HH clusters of forest loss were not correspondingly matched by HH clusters of fire points.

#### Spatial dependence between Forest loss and Fire density

The results of the diagnostic for spatial dependence indicated that a spatial error specification was suitable for the data of the 2001-2018 period (Table S2). Both the coefficient and the intercept estimates for each model were positive and significant in the spatial error models (*p* ≪ 0.01; Table 2). AIC value was lower in the spatial error model than that of its equivalent linear model. In addition, the Breusch-Pagan and Moran’s *I* tests showed no trace of heteroskedasticity and spatial autocorrelation of residuals, respectively.

**Table 2.**
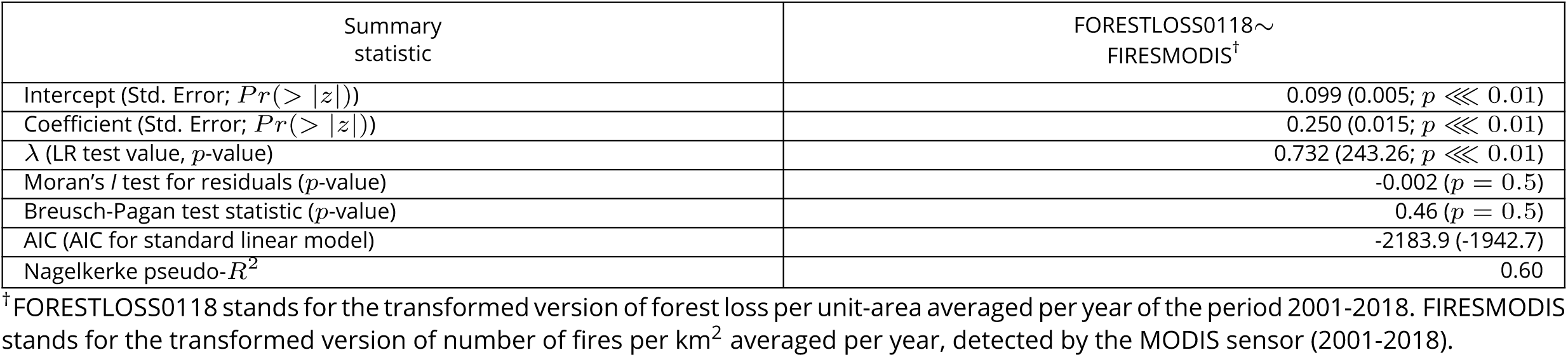
Spatial error model fitting results of forest loss as a function of fire density for the 2001-2018 period (MODIS fire data)

Lastly, considering only fire as a driver of forest loss, on average, each fire point detected by the MODIS sensor between 2001 and 2018 was associated with 1.5 ha forest loss, implying a substantial effect size of fire density on forest loss in the DR.

### Annual approach

#### Time series analysis

Using a paired *t*-test, I found no significant differences between proportional deforestation area originating from small and large clearings—*t=-2*.*08, df=17, p=0*.*053*. Furthermore, in several years of the study period (2001, 2003, 2011), the total area of deforestation originating from small clearings was greater than that from large clearings (see Fig. 6).

**Figure 6.**
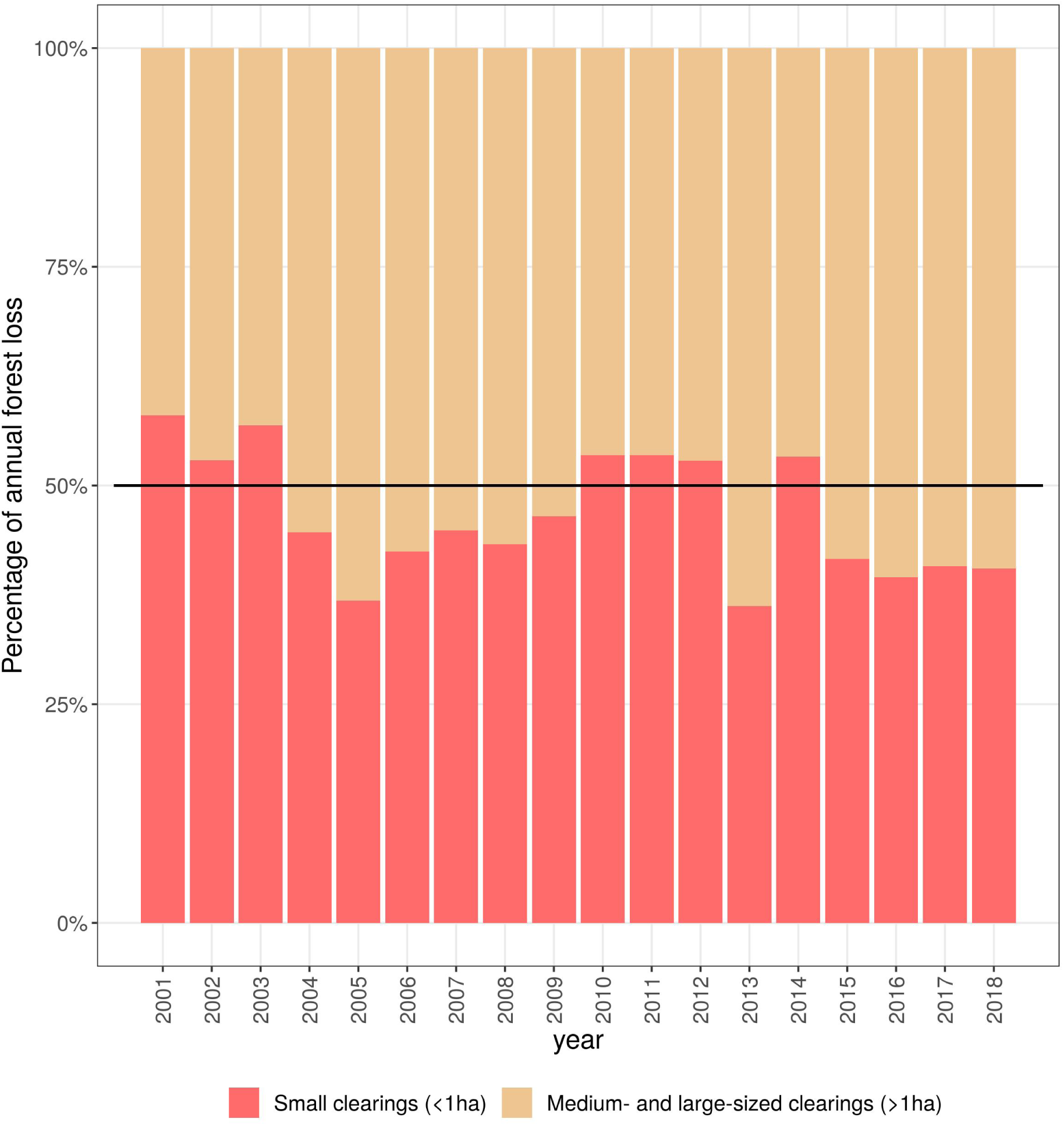
Composition of annual forest loss area by size of clearing

The yearly average forest loss area recorded in large clearings was 0.2 km^2^/100 km^2^, and reached a maximum of nearly 0.4 km^2^/100 km^2^. Further, the yearly average number of small clearings was 237 patches per 100 km^2^, and the maximum reached approximately 400 patches per 100 km^2^ (Figs. 7.A-B and S7.A-B). Regarding fire density, the MODIS sensor detected nearly two fire points per 100 km^2^ per year on average, and a maximum of 3.5 points per 100 km^2^ per year (Figs. 7.C and S7.C).

**Figure 7.**
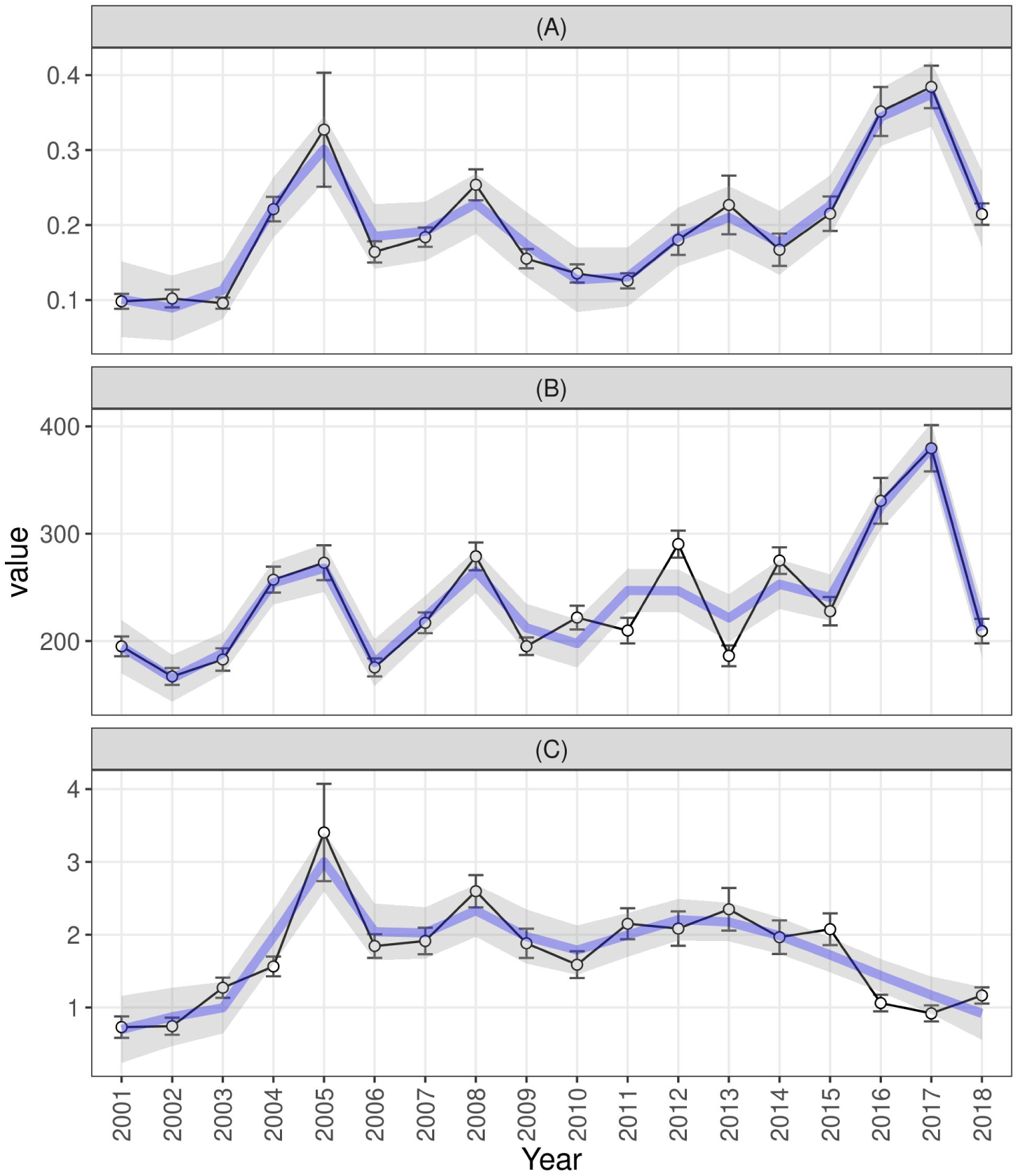
Yearly averages per 100 km^2^ of (A) Forest loss area (in km^2^) of large clearings (>1 ha in size); (B) Number of small clearings (<1 ha in size); (C) Number of fire points remotely sensed by the MODIS sensor in or around forest loss patches

Forest loss activity and fire occurrence showed a joint cyclical pattern of variation with a period of approximately 4 years from 2001 through 2013, with relative high peaks of activity in 2001, 2005, 2008 and 2012 (see Fig. S7). However, the time-series of forest loss and fire activity diverged considerably from each other, starting in 2014. In particular, forest loss increased rather steeply from 2014 to 2017, whereas the number of fire points decreased significantly during the same period (Fig. 7). Hence, this is the first time in the past two decades in which fire and forest loss followed diverging trends nationwide.

#### Spatio-temporal patterns

Regarding spatio-temporal features, both forest loss and fire density showed patterns of cyclical variation of their spatial autocorrelation, and featured multiple spatial layouts of HH clusters and LL clusters in shifting locations throughout the DR over the period under investigation. Moran’s *I* tests, which were applied to the transformed versions of the variables, yielded significant results for every year of the study period. In addition, the Moran’s *I* test statistic showed a cyclical and varied pattern for all the variables analyzed over the study period (Fig. 8).

**Figure 8.**
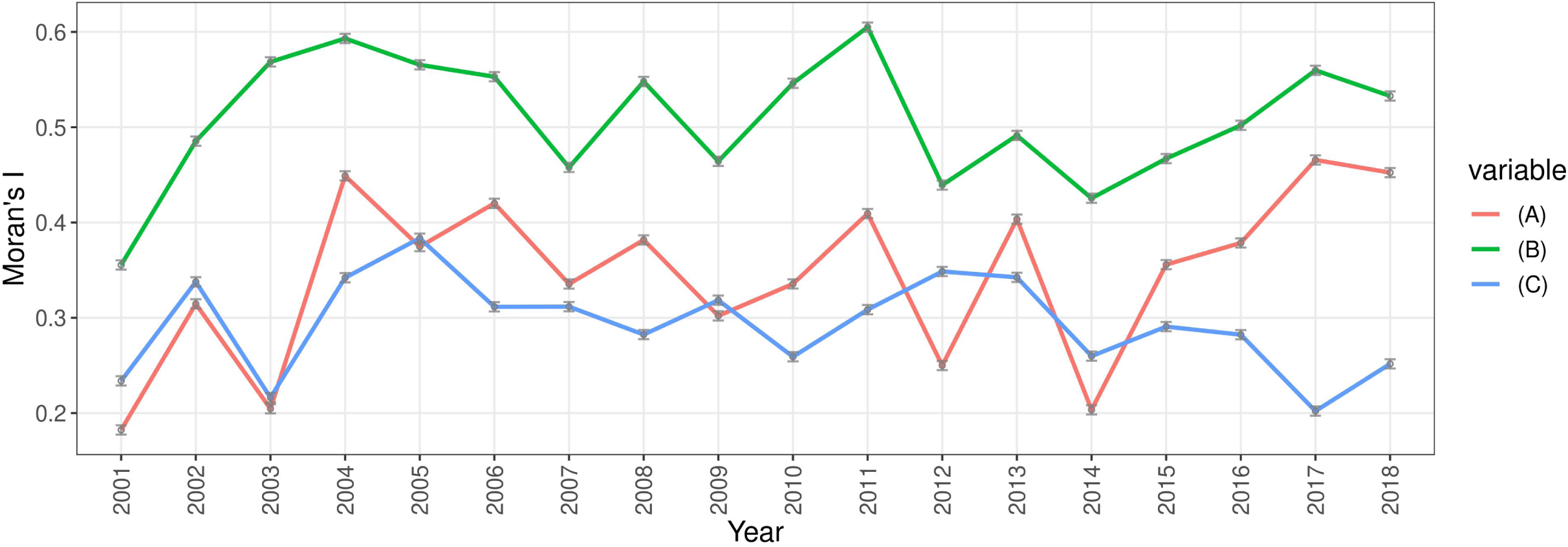
Moran’s *I* (used to assess spatial autocorrelation) evolution from 2001 to 2018 for the transformed versions of yearly averages per 100 km^2^ of: (A) Forest loss area of large clearings (>1 ha in size); (B) Number of small clearings (<1 ha in size); (C) Number of fire points located inside or around forest loss patches recorded by the MODIS sensor

Concerning patterns of forest loss, the HH clusters were concentrated mainly in five locations during the study period: Los Haitises-Samaná Peninsula, Cordillera Central, Sierra de Bahoruco, and Northwestern and Eastern Regions, the last being the largest tourism hub of the DR—including the resort town of Punta Cana and other tourist destinations (Figs. 9, 10, S8 and S9). The Eastern Region, in particular, showed very distinctive spatio-temporal patterns of forest loss over the study period; therefore, I analyzed that region separately.

**Figure 9.**
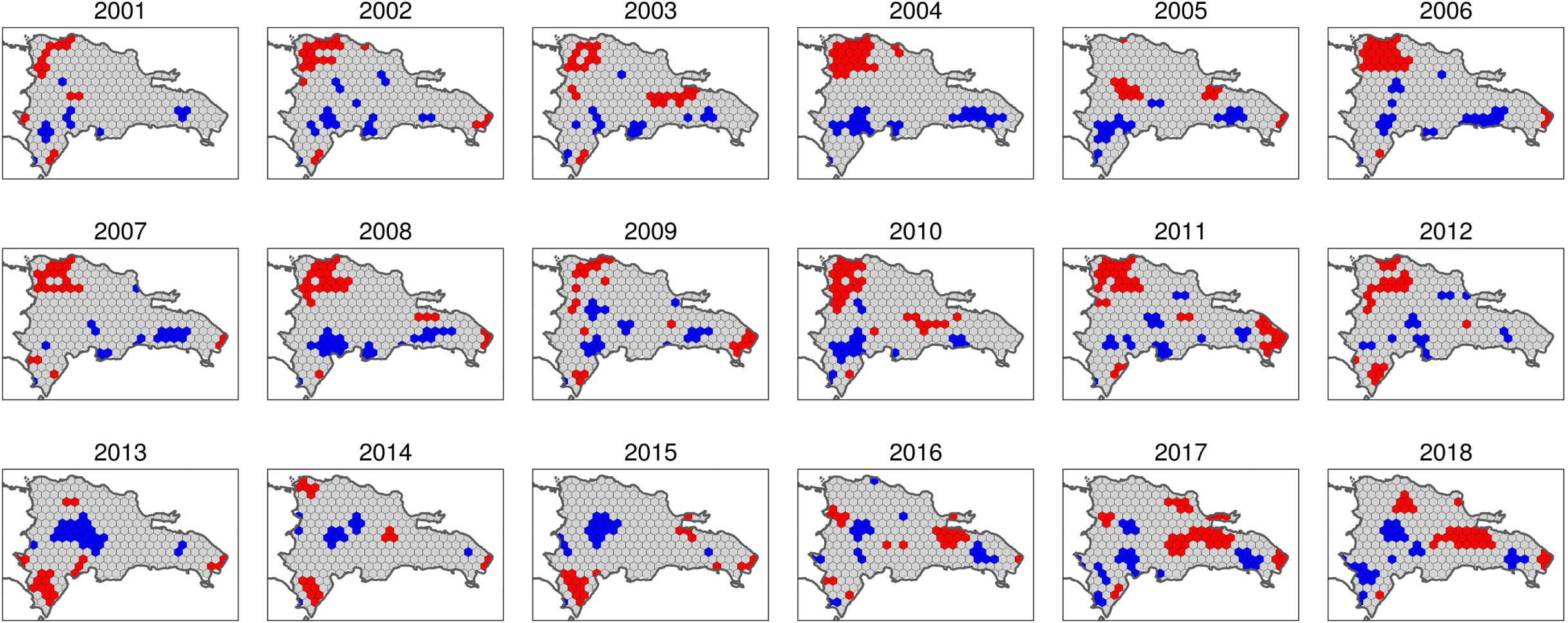
Yearly LISA maps (local indicators of spatial association maps) of transformed forest loss density data of large clearings. Red represents HH clusters, blue depicts LL clusters, and grey shows no significant spatial association.

**Figure 10.**
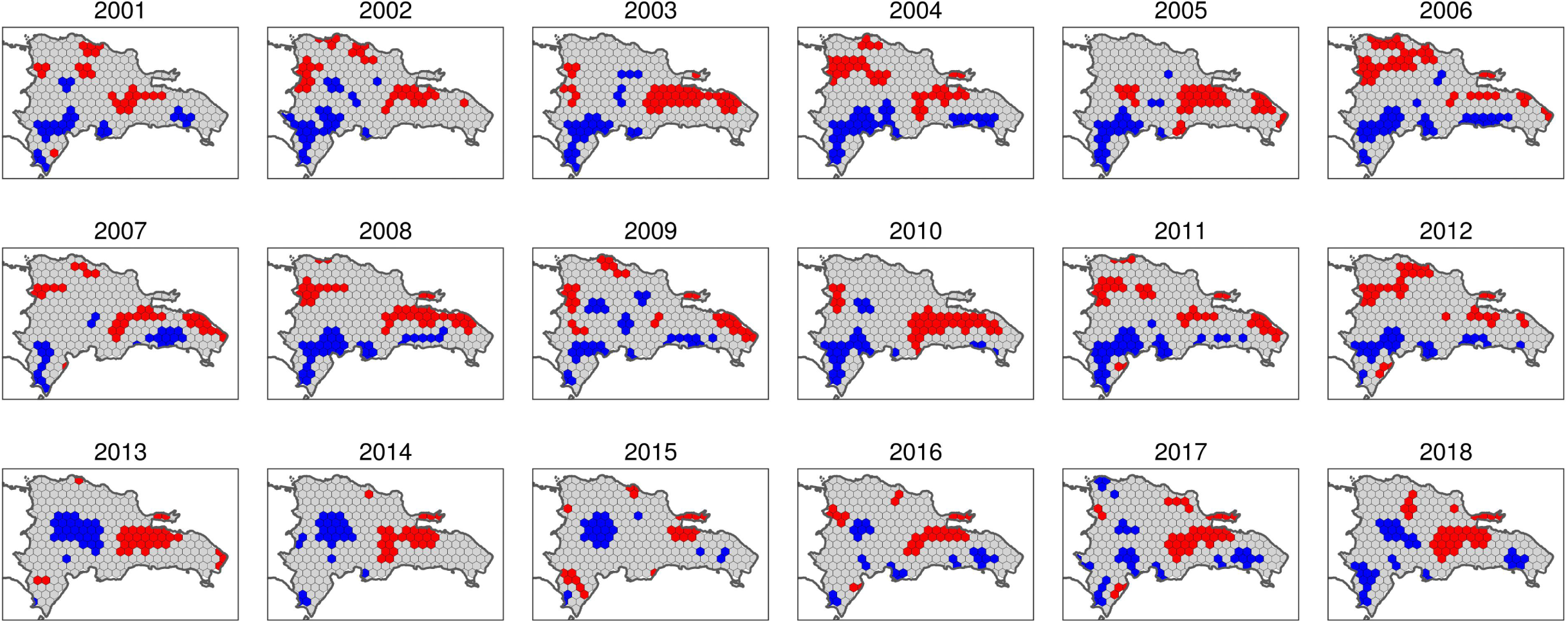
Yearly LISA maps (local indicators of spatial association maps) of transformed forest loss density data of small clearings. See Figure 9 for colour legend of hexagons.

During the three-year period 2001-2003, HH clusters of both small and large forest clearings were concentrated in Los Haitises-Samaná Peninsula and at the southern and northern ends of western DR. From 2004 to 2012, large forest clearings were significantly concentrated in the northwest of the DR—which peaked in 2004, and in the periods 2006-2008 and 2010-2011, in southern Cordillera Central and Sierra de Bahoruco. In addition, HH clusters of small clearings were widespread in Los Haitises in 2003, 2005, 2007-2008, and 2010. Subsequently, during the period 2013-2018, HH clusters of both large and small clearings were concentrated in Los Haitises and in Samaná Peninsula, as well as in western and southeastern portions of Cordillera Central and Sierra de Bahoruco. Of note was widespread deforestation in Los Haitises and its southern end, which comprises an active oil palm plantation. These hotspots are shown on the LISA maps by large HH clusters of small clearings during the period 2013-2014, as well as by HH clusters of both large and small clearings in the period 2017-2018.

In addition, concerning the Eastern Region, HH clusters of large clearings began to develop in 2002 and stopped in subsequent years, then emerged intermittently from 2005 onwards, showing peaks of activity in 2009 and 2011 and a steady increase between 2013 and 2018. Notably, HH clusters of small clearings were detected in this region in 2003, in 2005-2009, and in years 2011 and 2013, but no new clusters of this type were observed in subsequent years.

Regarding fire density, during the entire period investigated, HH clusters were concentrated especially in the western half of the DR, particularly in the Northwestern Region, Sierra de Bahoruco, and Cordillera Central (Figs. 11 and S10). Also, during both the first years and in the middle of the period, HH clusters were present in Los Haitises and Samaná Peninsula.

**Figure 11.**
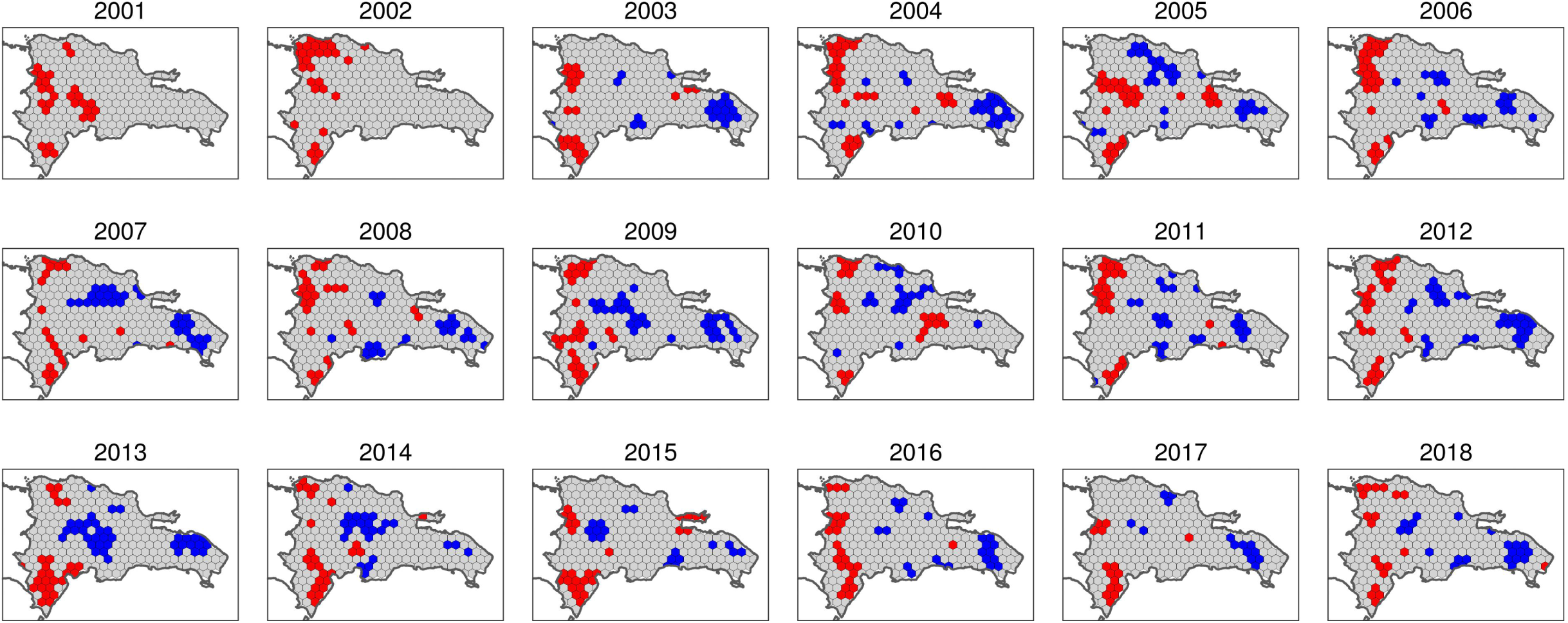
Yearly LISA maps (local indicators of spatial association maps) of transformed MODIS fire density data. See Figure 9 for colour legend of hexagons.

**Figure 12.**
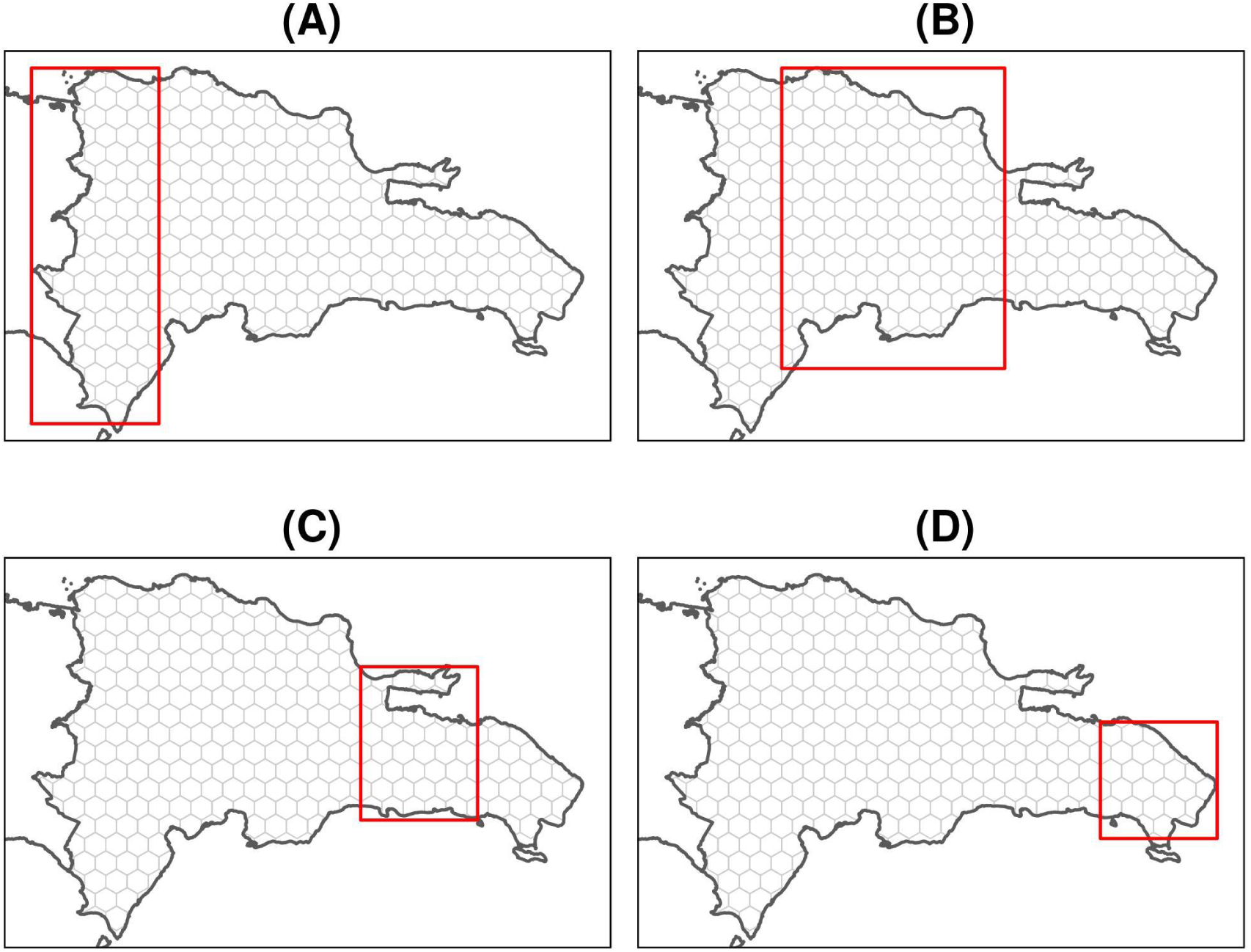
Regions for annual model analyses. (A) Western, (B) Central, (C) Los Haitises-Samaná, and (D) Eastern.

As shown in the LISA maps of MODIS fire density, the spatial patterns of fire density slightly resembled those of forest loss over the period under study (Fig. 11). However, the degree of agreement between forest loss and fire density was greater in the Western and Central Regions—Sierra de Bahoruco, Northwestern Region, Cordillera Central—than in the eastern half of the country—Los Haitises and the Eastern Region. Particularly, although the eastern half showed extensive forest loss activity, few HH clusters of fire density were recorded in this region during the period under investigation. In fact, during the six-year period 2013-2018, HH clusters of fire completely disappeared from the Eastern Region (Fig. 11). Hence, fire activity showed a diverging trend in relation to that of deforestation in Los Haitises and the Eastern Region.

In addition, three remarkable features regarding the distribution of HH clusters of fire density merit mention in this section. (Figs. 11 and S10). The first is a large concentration of HH clusters in 2005 over southern Cordillera Central, related to an uncontrolled wildfire that devastated almost 80 km^2^ of pine forest. As a result, more than 100 fire points per 100 km^2^ were reached, which is a historical record. Second, for three years in a row—2013, 2014, 2015—the MODIS sensor detected a high concentration of hotspots over Sierra de Bahoruco, attributable to multiple wildfires that swept large areas of different types of mountain forests during those years. Third, in 2014 and 2015, the MODIS sensor detected a relatively high number of fire points in Valle Nuevo, southern Cordillera Central, which are depicted in Fig. S10 as HH clusters, and which are also consistent with the fire history of the area.

Finally, the spatial error models yielded consistent results for forest loss as a function of fire density (Table 3). The main finding was that, when modeling the variables over the entire grid—i.e., nationwide analysis— fire density was significantly associated with forest loss, which is consistent with the results of the long-term approach. Particularly, both fire density coefficient and intercept were significant in every annual model, regardless of the size of deforestation clearings, whether large or small. Moreover, regional subsets showed that fire density was a suitable predictor of forest loss most of the time in Western and Central regions, whereas in Los Haitises-Samaná Peninsula and the easternmost region, fire density failed as a predictor of forest loss for many years.

**Table 3.** Number of years in which the coefficients of the annual spatial error models were not significant, considering the entire grid and different regional subsets.

| Variables of the models (transformed versions) | Regional subset (see Fig. 12) | Number of years<sup>†</sup> (listing in parenthesis) |
| --- | --- | --- |
| Forest-loss per unit-area in large clearings vs. MODIS fire density | Entire grid | - |
|  | Western | - |
|  | Central | 1 year (16) |
|  | Los Haitises-Samaná | 11 years (1, 2, 7, 9-14, 16, 17) |
|  | Eastern | 9 years (1-4, 7, 10, 15-17) |
| Forest-loss per unit-area in small clearings vs. MODIS fire density | Entire grid | - |
|  | Western | - |
|  | Central | 3 years (4, 12, 16) |
|  | Los Haitises-Samaná | 7 years (1, 2, 4, 7, 12, 14, 17) |
|  | Eastern | 15 years (1-12, 15-17) |

<sup>†</sup>Number of years with non-significant coefficient at $\alpha = 0.01$

## Discussion

I hypothesized that fire and forest loss were significantly associated during the first 18 years of the 21st Century in the DR, and that fire was a suitable predictor of forest loss, regardless of the size of the forest clearings. The evidence found in the present study supports this hypothesis consistent with other studies that found a significant association between forest loss and slash-and-burn agriculture (Lloyd and León, 2019; Myers et al., 2004; Wendell Werge, 1974; Zweifler et al., 1994). Moreover, the association between fire and forest loss is particularly consistent in the western half of the DR, which is likely due to the more pronounced dry season in that region and to the presence of large mountain systems, i.e., Cordillera Central, where shifting agriculture is widespread.

However, the evidence also suggests that, in the eastern half of the country, which includes Los Haitises and the easternmost region, fire was not a suitable predictor of forest loss. Two conjectures may explain this finding: 1) Frequent cloudy skies over the region, which may prevent the MODIS sensor from recording fire hotspots; 2) Factors other than fire that may drive forest loss, such as commodity-driven agriculture, shifting agriculture by means of downing vegetation without burning—or by indeed performing burns but with little impact on forest cover—and expansion of tourism infrastructure facilities. The first conjecture is unlikely to explain the observed pattern, since fire activity in cloudy conditions, considered on an annual average basis, would have little effect as a driver of pervasive deforestation. The second conjecture provides a more likely explanation for deforestation peaks not associated with fires, since it fits quite well with the tree cover decimation mechanisms that are typically used in this part of the DR, i.e., forest clearing to expand shifting agriculture driven primarily by subsistence needs. Since there are many contextual differences between Los Haitises and the easternmost tourism hub, I discuss the implications of holding this hypothesis true for each area separately.

In Los Haitises National Park, shifting agriculture was likely the most suitable driver of deforestation, since it is a well-documented concern in this protected area (Dirección Nacional de Parques, 1991; Gesto de Jesús, 2016). Shifting agriculture is commonly driven by slash-and-burn systems, but in this case the “burn” component was likely to have little effect as a driver of deforestation in that area. Overall, the evidence suggests that shifting agriculture was widespread within the protected area, particularly in the period 2014-2017. However, the political and socioeconomic circumstances that led to a deforestation peak in Los Haitises and surroundings remain unknown. Future research may provide insights into the specific causes that explain this peak in shifting agriculture, and may also provide guidelines on how to prevent the recurrence of deforestation peaks in the future, given that Los Haitises is an important protected area of the country.

It is worth mentioning that, in this part of the DR, another probable source of deforestation without burning is the frequent renewal by cutting of palm trees in a large plantation situated just south of Los Haitises—a typical case of commodity-driven deforestation. Although this plantation is outside the boundaries of the national park, its impact on the biodiversity and ecology of the area is unknown.

Finally, in the easternmost region, much of the forest loss activity was probably driven by the expansion of tourism facilities, and by increased agricultural and livestock activities, ultimately caused by a higher demand from tourism. This is a concerning trend for the future of the DR forests, because although the protected areas of the region are relatively well preserved, there is a lack of policies aimed at the conservation and proper management of the forests in the vicinity of tourism facilities.

Regarding spatial patterns, I also hypothesized that both forest loss and fire experienced a growing spatial autocorrelation over the study period. Although a high degree of spatial autocorrelation was a common characteristic in both forest loss and fire density variables over the study period, no evidence was found to support a hypothesis of a growing autocorrelation trend. Instead, a cyclical variation of autocorrelation was the most common feature observed, which I interpret as a consequence of both deforestation recovery and drought-no drought cycles. However, further research is needed to determine the precise causes of those singular cycles.

I also suggest that the results of this study may assist decision makers in designing effective policies to prevent wildfires and forest loss. More specifically, I recommend focusing on forest loss and fire prevention in the core zones of protected areas, and implementing a natural regeneration program by letting nature evolve on its own where forest cover is lacking, especially in the buffer zones of mountain protected areas of western DR. Of particular concern are the core and buffer zones of José del Carmen Ramírez, Los Haitises, Valle Nuevo and Sierra de Bahoruco National Parks, as well as other protected areas in the northern margin of Cordillera Central. These areas, especially the hotspot locations highlighted as HH clusters in the LISA maps, require more attention and resources to prevent wildfires and deforestation. In addition, there is a lack of special policies to prevent deforestation in areas surrounding major cities and tourism hubs. The recommendations in these areas are to avoid further deforestation activities and to allow regeneration in the most affected ecosystems, such as wetlands and areas of high endemism that previously had forests.

The main limitations of this study were those imposed by the intrinsic characteristics of the data available, which are ultimately related to the data acquisition mechanisms of the MODIS optical sensor. Although detecting fires under cloud cover is virtually impossible with this sensor, I surmise that the impact of false negatives on yearly analyses is quite limited. Another constraint met in this study was the use of fixed-size cells for the computations of zonal statistics, which may have prevented the determination of multiscalar patterns. Therefore, future research using regular and non-regular grids as zone layers, or taking advantage of computer vision and machine learning techniques, may provide insights about the significant multiscalar association patterns that may exist between forest loss and fire.

In conclusion, fire is a fairly common feature associated with shifting agriculture, so assessing the former is an indirect means for understanding the latter, which ultimately may help prevent future impact on forest ecosystems. Therefore, proper fire assessment using remotely collected data and advanced spatial statistical techniques may inform land management policies and conservation strategies to help reduce forest loss, particularly in protected areas, mountain areas, and the vicinity of tourism hubs. The analytical approaches used and the results obtained in this study hold potential to assist in this task.

## Acknowledgements

The author acknowledges the use of data and imagery from LANCE FIRMS operated by NASA’s Earth Science Data and Information System (ESDIS) with funding provided by NASA Headquarters.

Version 4 of this preprint has been peer-reviewed and recommended by Peer Community In Forest and Wood Sciences (https://doi.org/10.24072/pci.forestwoodsci.100005)

## Fundings

This research received no specific grant from any funding agency in the public, commercial, or not-for-profit sectors. This work was supported by Universidad Autónoma de Santo Domingo.

## Conflict of interest disclosure

The author of this article declare that he has no financial conflict of interest with the content of this article.

## Data, script and code availability

The data that support the findings of this study are openly available in Zenodo at https://doi.org/10.5281/zenodo.5682104. The scripts used for data curation, analysis and visualisation are available in Zenodo at https://doi.org/10.5281/zenodo.6762306.

## Supplementary Information

### Supplementary methods

To generate “the noise-free versions of the FIRMS collections”, I wrote an algorithm that spotted extremely dense point clusters. Afterward, I confirmed whether those clusters fell into industrial or landfill areas, by visually checking with base maps and satellite images. In most cases, those points were tagged as “other static land source” in the “Type” field of the datasets. Points that met at least the visual examination criteria were excluded from the dataset. In addition, I excluded all points tagged with a low confidence value (Fig. S1).

In addition, I applied a mask comprising the DR land area to each dataset used in the study. I generated the mask by combining a shapefile containing the international DR border, downloaded from https://www.one.gob.do/informaciones-cartograficas/shapefiles, with the datamask included in the forest change dataset. Permanent water bodies were excluded from the analysis, using their extent area as seen in the 2000 Landsat ETM+ imagery.

**Figure S1.**
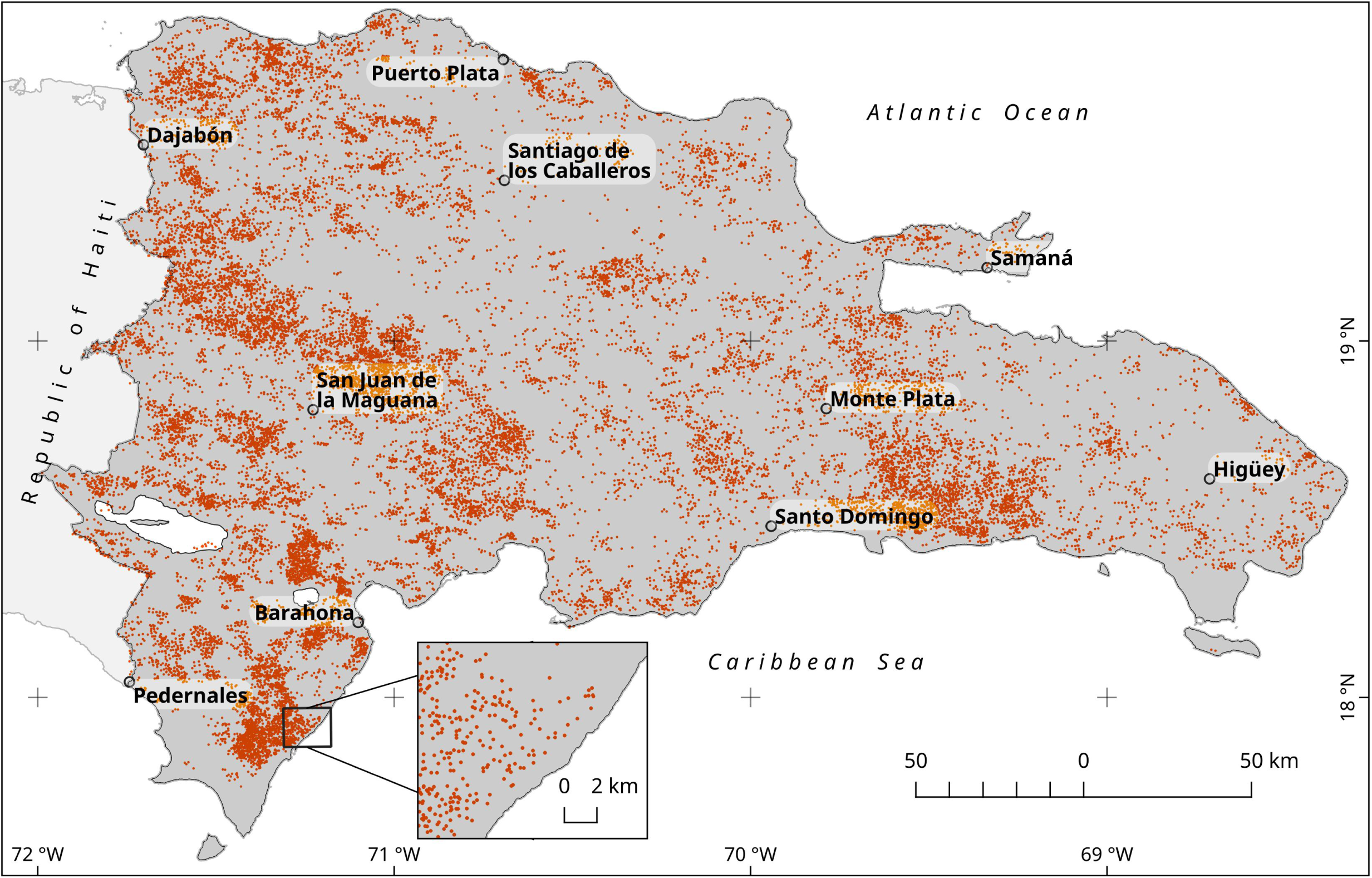
MODIS fire points/hotspots from 2001 to 2018 for the DR mainland. This is a noise-free version of the original dataset, which excludes unrelated fire points (e.g., burning landfills and industrial furnaces). See text for details.

## Supplementary data for the results section

### MODIS data consistency and sensitivity assessment

**Figure S2.**
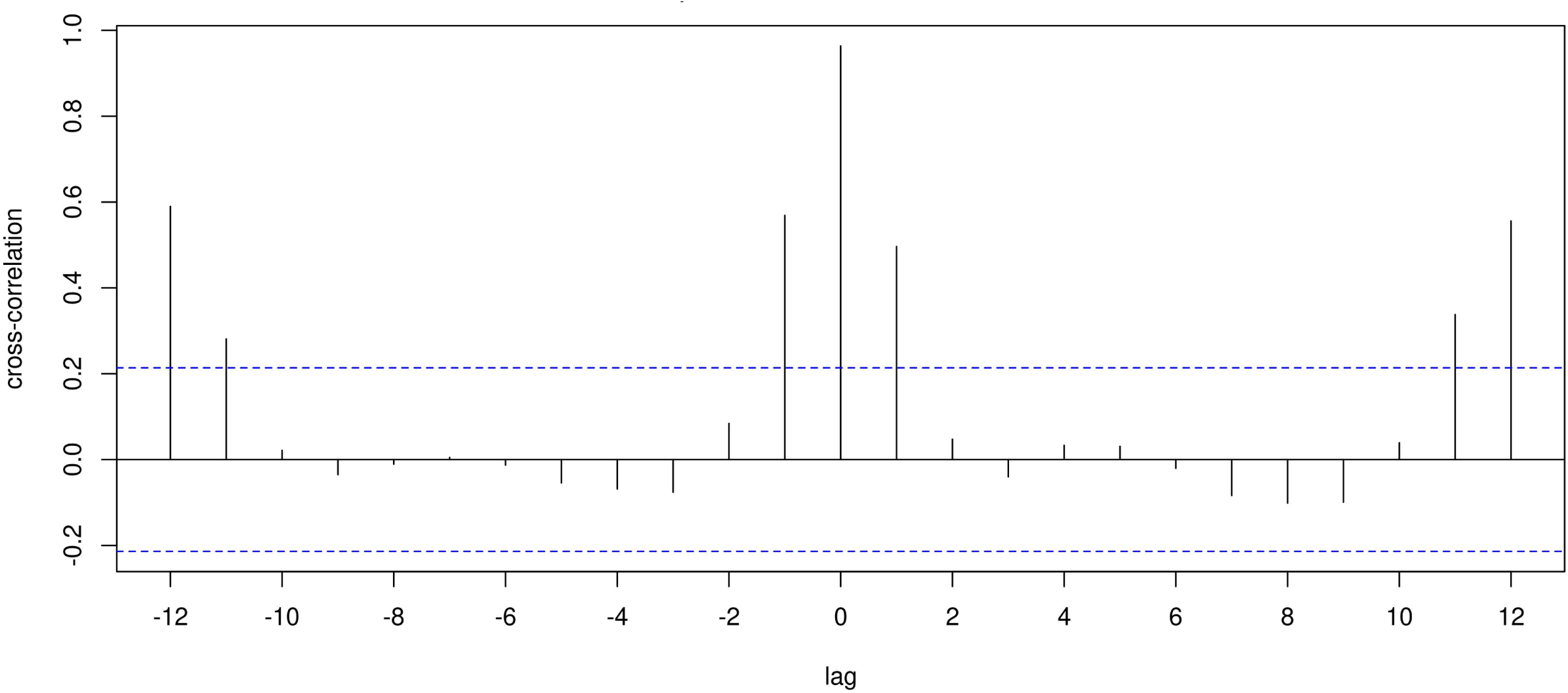
Cross-correlation of number of fire points per month sensed by MODIS and VIIRS sensors for the period 2012-2018

**Figure S3.**
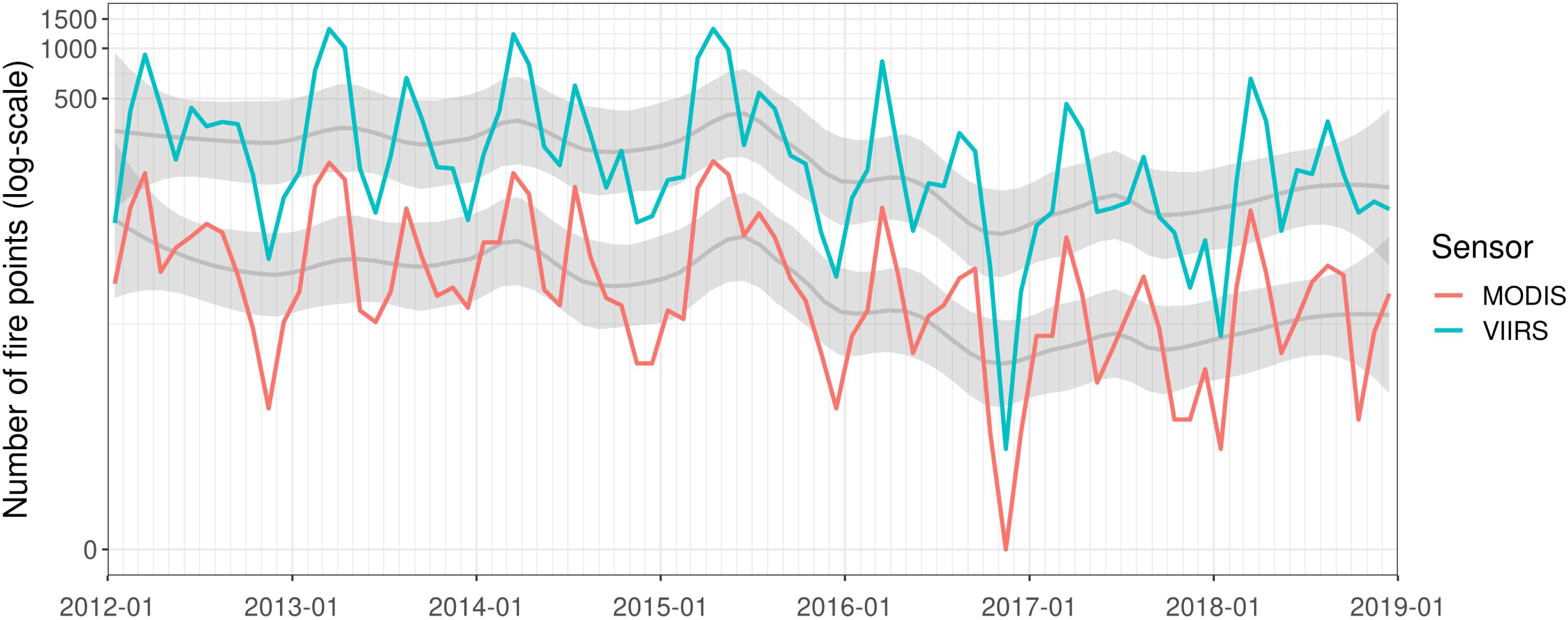
Number of fire points per month sensed by MODIS and VIIRS sensors for the period 2012-2018

**Figure S4.**
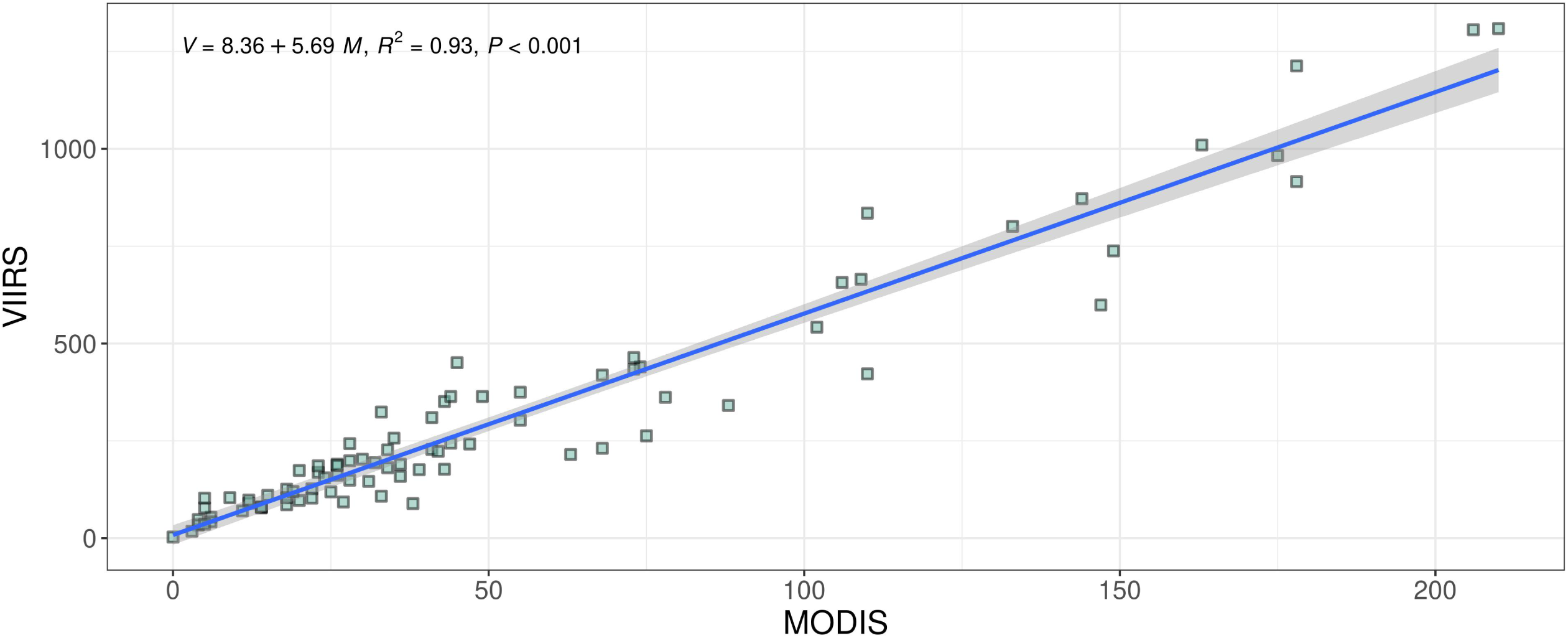
Scatter plot of the number of fire points per month sensed by MODIS and VIIRS sensors for the period 2012-2018

### Long-term approach

**Table S1.** Transformation parameters and normality test results for forest loss and fire variables

| Variable | Tukey's Ladder of Powers, $\lambda$ | Shapiro-Wilk test,<br>$W$ ($p$-value) | Moran's $I$ test,<br>$I$ ($p$-value) |
| --- | --- | --- | --- |
| Average forest loss per unit area per year (2001-2018) | $\lambda = 0.33$ | $W = 0.99$ ($p = 0.81$) | $I = 0.48$ ($p \lll 0.01$) |
| Average fire density per km<sup>2</sup> per year (MODIS dataset) (2001-2018) | $\lambda = 0.33$ | $W = 0.98$ ($p < 0.01$) | $I = 0.55$ ($p \lll 0.01$) |

**Figure S5.**
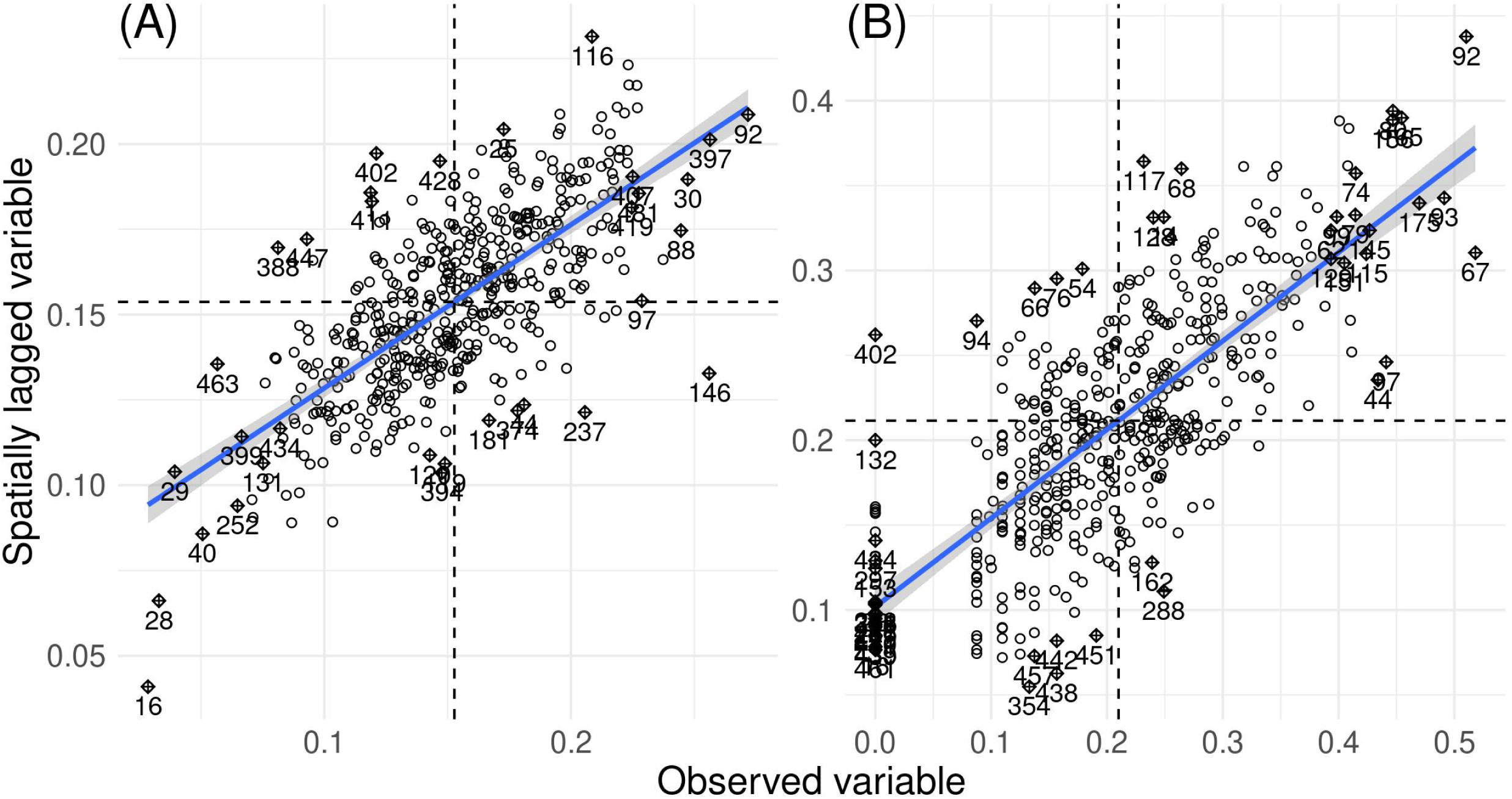
Moran scatterplots of the transformed versions of the analyzed variables. (A) Average forest loss per unit area per year of the period 2001-2018. (B) average number of fire points per km^2^ per year in the period 2001-2018 using MODIS dataset.

**Table S2.** Lagrange Multiplier tests for spatial dependence in linear regression models of forest loss as a function of fire density for the period 2001-2018 (MODIS fire data)

|  | FORESTLOSS0118 ~ FIRESMODIS<sup>†</sup> |  |
| --- | --- | --- |
| Lagrange Multiplier test | Statistic | $p$ value |
| For error dependence (LMerr) | 330.00 | $\lll 0.01$ |
| For a missing spatially lagged dependent variable (LMlag) | 227.22 | $\lll 0.01$ |
| Robust variant of LMerr | 106.49 | $\lll 0.01$ |
| Robust variant of LMlag | 3.72 | 0.05 |

<sup>†</sup>FORESTLOSS0118 stands for the transformed version of forest loss per unit-area averaged per year of the period 2001-2018. FIRESMODIS stands for the transformed version of number of fires per km<sup>2</sup> averaged per year, detected by the MODIS sensor (2001-2018)

### Annual approach

**Figure S6.**
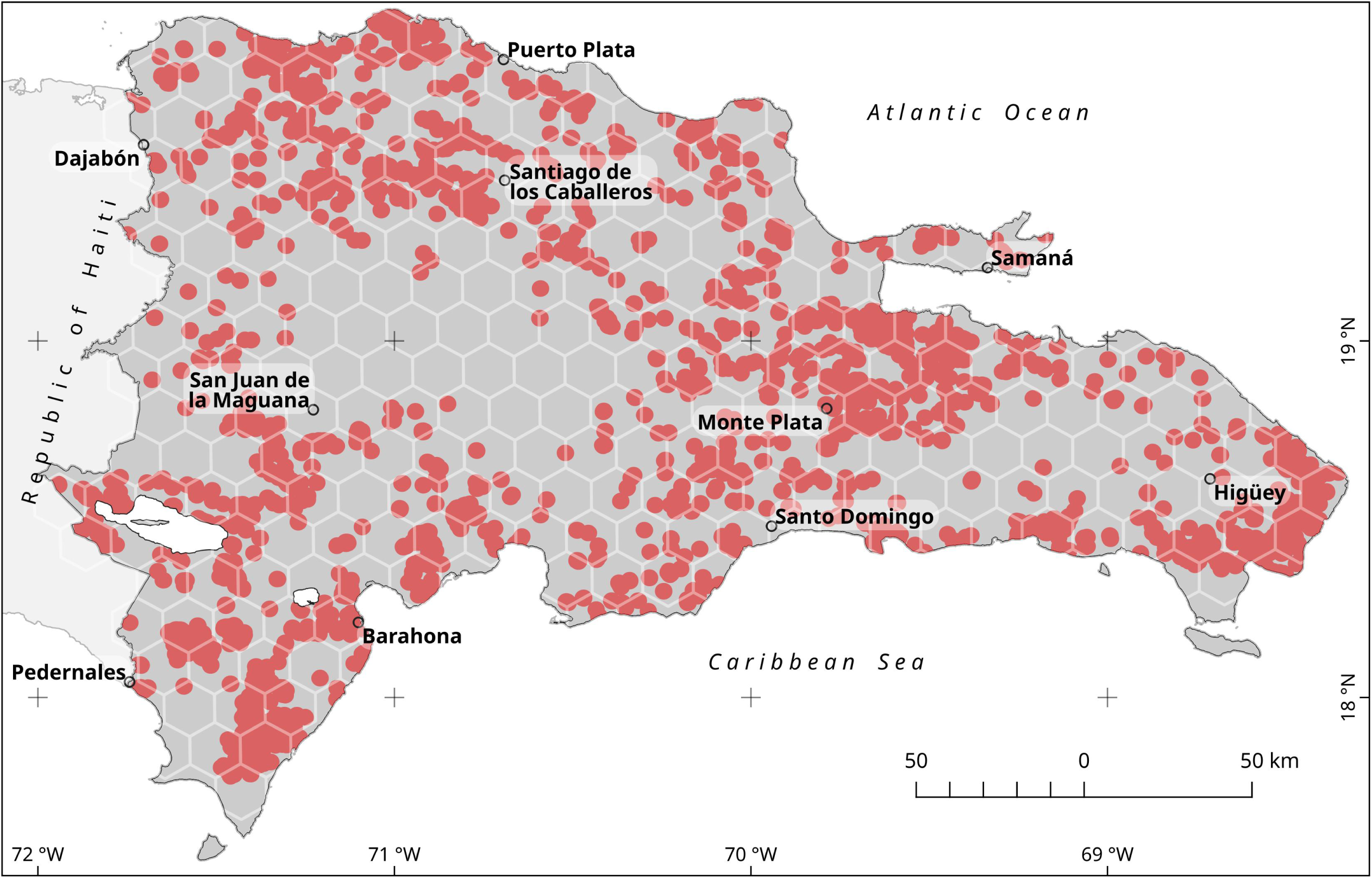
Example of the 2013 forest loss areas and their vicinity (red shaded areas) used in the annual trend approach. These areas were generated by adding a buffer zone of 2.5 km around each patch larger than 1 ha in area from the loss year dataset (MC Hansen et al., 2013). The hexagonal grid, depicted as an overlay, was used for zonal statistics computations. See text for details.

**Figure S7.**
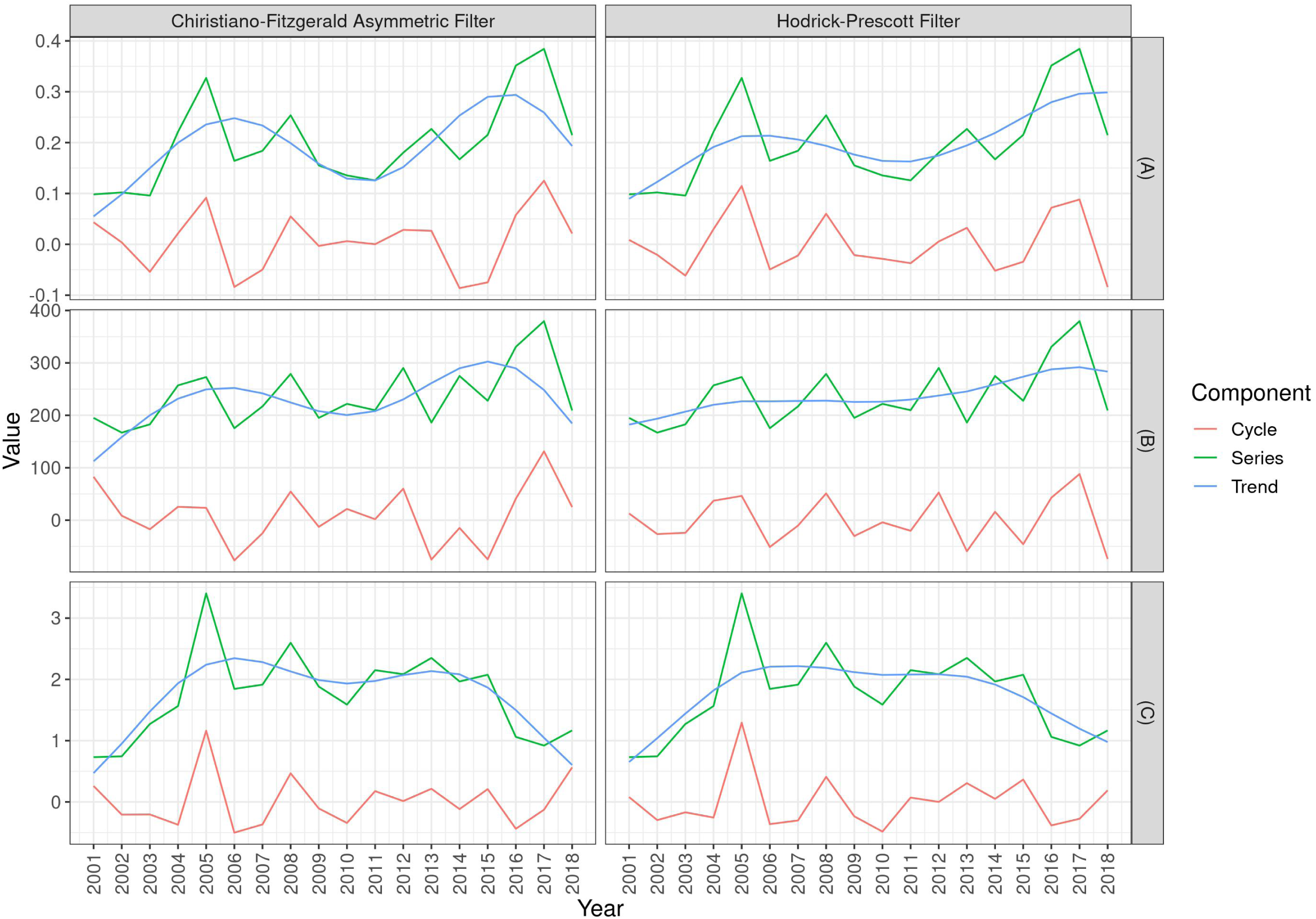
Time series decomposition of yearly averages per 100 km^2^ of (A) Forest loss area (in km^2^) of large clearings (>1 ha in size); (B) Number of small clearings (<1 ha in size); (C) Number of fire points remotely sensed by the MODIS sensor in or around forest loss patches

**Figure S8.**
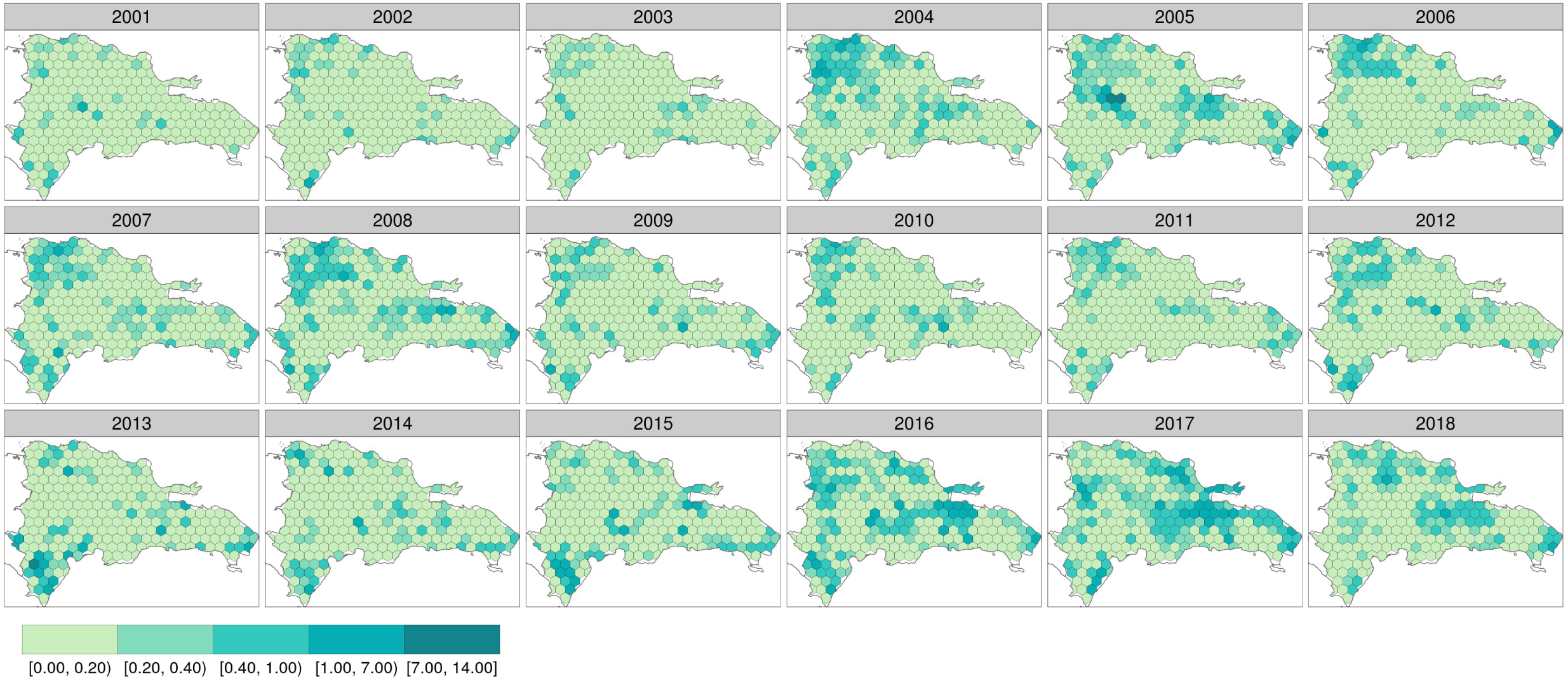
Yearly forest loss area (in km^2^ per 100 km^2^) from patches greater than 1 ha in size for the period 2001-2018

**Figure S9.**
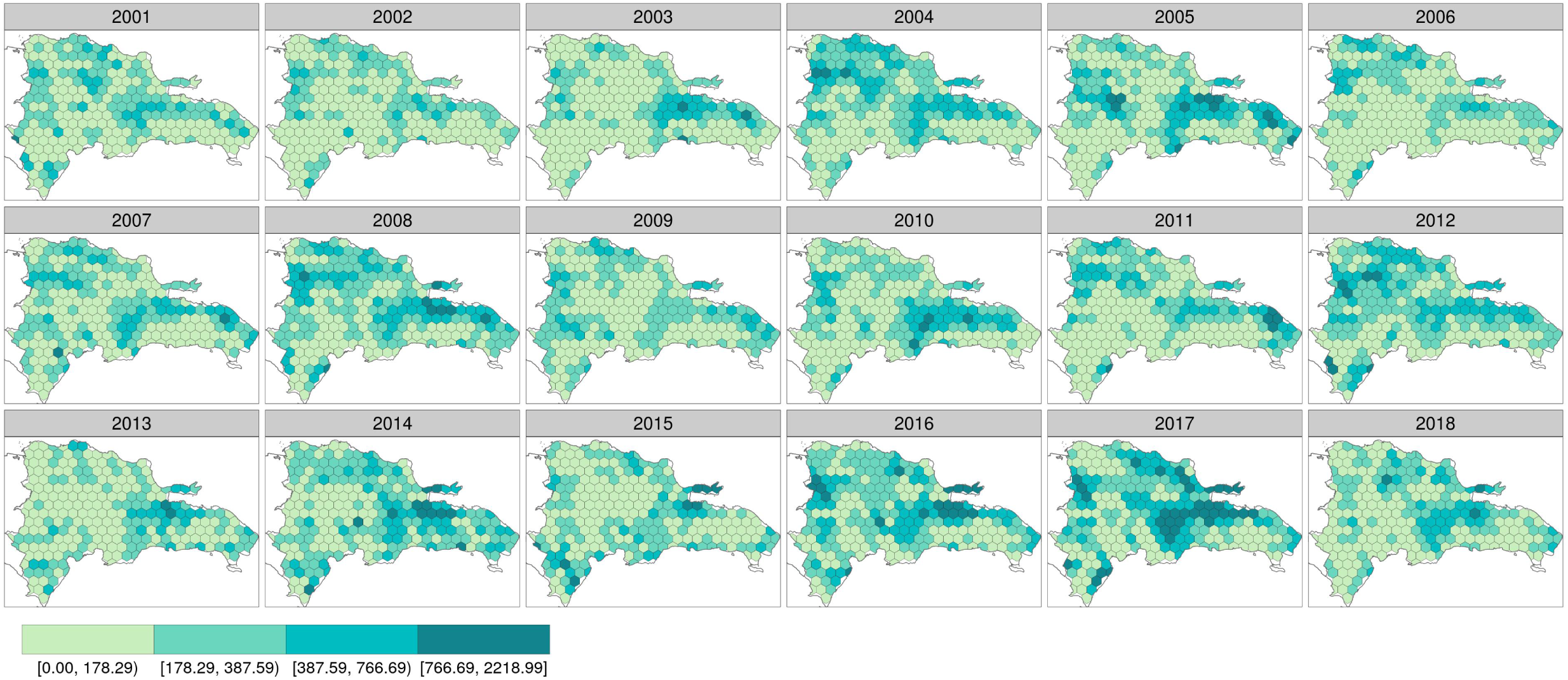
Yearly number of forest loss patches smaller than 1 ha (in km^2^ per 100 km^2^) for the period 2001-2018

**Figure S10.**
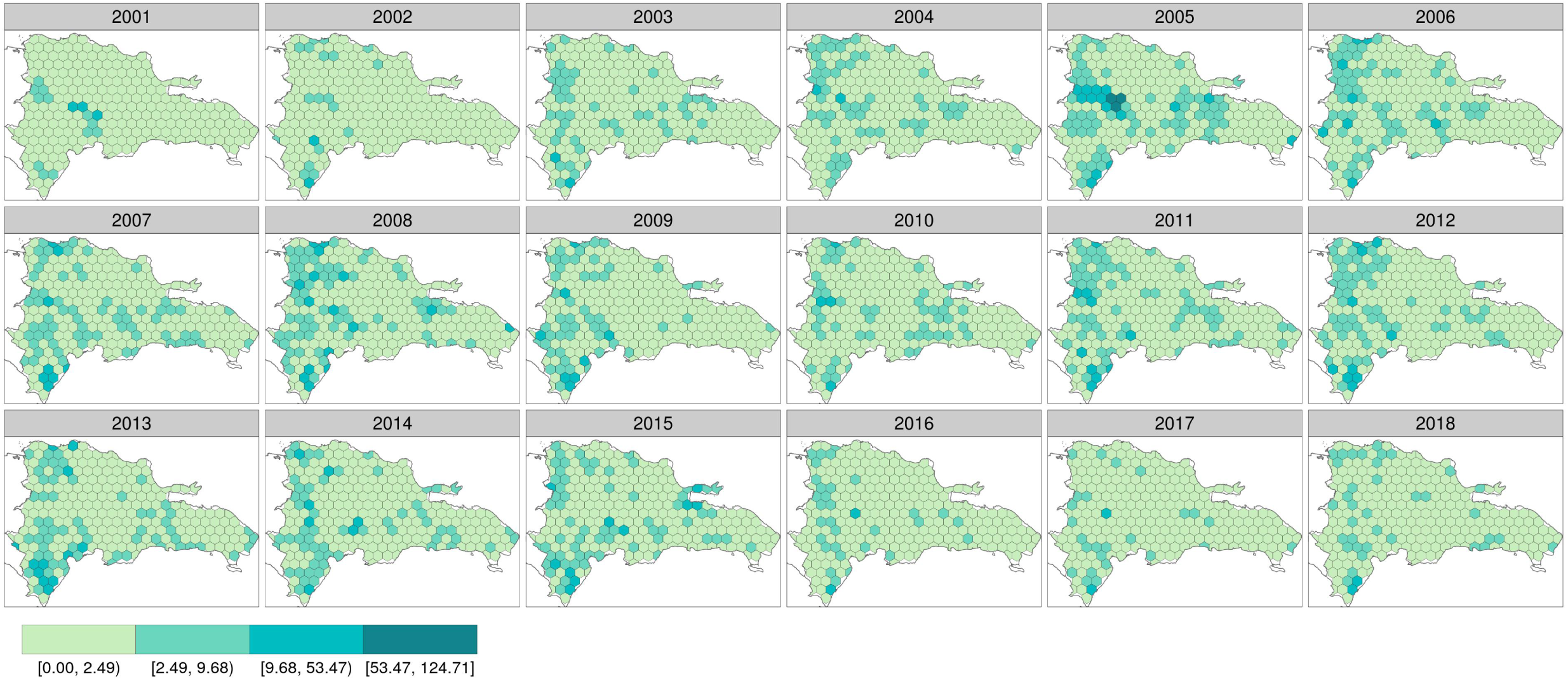
Yearly number of MODIS fire points per 100 km^2^ within patches of forest loss and surroundings for the period 2001-2018

## Notes

https://doi.org/10.5281/zenodo.6762306

https://doi.org/10.5281/zenodo.5682104

### Competing Interest Statement

The authors have declared no competing interest.

### Summary of Updates

Revised version after round #2

## References

Anselin L (1995). Local indicators of spatial association—LISA. Geographical analysis 27, 93–115. https://doi.org/10.1111/j.1538-4632.1995.tb00338.x.

Anselin L (1996). The Moran scatterplot as an ESDA tool to assess local instability in spatial association. In: Spatial Analytical Perspectives on GIS in Environmental and Socio-Economic Sciences. Ed. by Fischer M, Scholten H, and Unwin D. Taylor and Francis. Chap. 8, pp. 111–125. https://doi.org/10.1201/9780203739051.

Anselin L (2013). Spatial econometrics: methods and models. Vol. 4. Springer Science & Business Media. https://doi.org/10.1007/978-94-015-7799-1.

Anselin L and SJ Rey (2010). Perspectives on spatial data analysis. In: Perspectives on Spatial Data Analysis. Ed. by Anselin L and Rey SJ. Berling, Heidelberg: Springer. Chap. 1, pp. 1–20. https://doi.org/10.1007/978-3-642-01976-0.

Balcilar M (2019). mFilter: Miscellaneous Time Series Filters. Available on-line at https://CRAN.R-project.org/package=mFilter.

Bivand R, M Altman, L Anselin, R Assunção, and O Berke (2017). Package ‘spdep’. Available on-line at https://cran.r-project.org/web/packages/spdep/index.html.

Bivand R, J Hauke, and T Kossowski (2013). Computing the Jacobian in Gaussian spatial autoregressive models: An illustrated comparison of available methods. Geographical Analysis 45, 150–179. https://doi.org/10.1111/gean.12008.

Bivand R and G Piras (2015). Comparing Implementations of Estimation Methods for Spatial Econometrics. Journal of Statistical Software 63, 1–36. https://doi.org/10.18637/jss.v063.i18.

Bivand R and DWS Wong (2018). Comparing implementations of global and local indicators of spatial association. TEST 27, 716–748. https://doi.org/10.1007/s11749-018-0599-x.

Bivand RS, E Pebesma, and V Gomez-Rubio (2013). Applied spatial data analysis with R, Second edition. Springer, NY. https://doi.org/10.1007/978-1-4614-7618-4.

Breusch TS and AR Pagan (1979). A simple test for heteroscedasticity and random coefficient variation. Econometrica: Journal of the Econometric Society, 1287–1294. https://doi.org/10.2307/1911963.

Buřivalová Z, SJ Hart, VC Radeloff, and U Srinivasan (2021). Early warning sign of forest loss in protected areas. Current Biology 31, 4620–4626. https://doi.org/10.1016/j.cub.2021.07.072.

Cámara Artigas R (1997). República Dominicana: Dinámica del medio físico en la región Caribe (Geografía Física, sabanas y litoral). Aportación al conocimiento de la tropicalidad insular. PhD thesis. Departamento de Geografía Física y Análisis Geográfico Regional, Facultad de Geografía e Historia, Universidad de Sevilla. https://doi.org/https://hdl.handle.net/11441/85112.

Cano E and A Veloz (2012). Contribution to the knowledge of the plant communities of the Caribbean-Cibensean Sector in the Dominican Republic. Acta Botanica Gallica 159, 201–210. https://doi.org/10.1080/12538078.2012.696933.

Curtis PG, CM Slay, NL Harris, A Tyukavina, and MC Hansen (2018). Classifying drivers of global forest loss. Science 361, 1108–1111. https://doi.org/10.1126/science.aau3445.

Department of Economic and Social Affairs of the United Nations Secretariat (2009). The millennium development goals report 2009. https://doi.org/10.18356/00399789-en.

Dirección Nacional de Parques (1991). Plan de uso y gestión del parque nacional Los Haitises y áreas periféricas. Dominican Republic Santo Domingo: Agencia Española de Cooperación Internacional y Agencia de Medio Ambiente de la Junta de Andalucía, pp. 1–381.

Gesto de Jesús EM (2016). Motores de deforestación en el Parque Nacional Los Haitises y uso de hábitat de anidación del Gavilán de la Española (Buteo ridgwayi), República Dominicana. Available on-line at http://repositorio.bibliotecaorton.catie.ac.cr/handle/11554/8601. MA thesis. CATIE, Turrialba (Costa Rica).

Greenberg JA and M Mattiuzzi (2018). gdalUtils: Wrappers for the Geospatial Data Abstraction Library (GDAL) Utilities. Available on-line at https://CRAN.R-project.org/package=gdalUtils.

Hager J and TA Zanoni (1993). La vegetación natural de la República Dominicana: una nueva clasificación. Moscosoa 7. Available on-line at https://www.biodiversitylibrary.org/item/181315, 39–81.

Hák T, S Janoušková, and B Moldan (2016). Sustainable Development Goals: A need for relevant indicators. Ecological Indicators 60, 565–573. ISSN: 1470-160X. https://doi.org/10.1016/j.ecolind.2015.08.003.

Hansen M, P Potapov, B Margono, S Stehman, S Turubanova, and A Tyukavina (2014). Response to Comment on “High-resolution global maps of 21st-century forest cover change”. Science 344, 981–981. https://doi.org/10.1126/science.1248817.

Hansen MC, PV Potapov, R Moore, M Hancher, SA Turubanova, A Tyukavina, D Thau, S Stehman, SJ Goetz, TR Loveland, et al. (2013). High-resolution global maps of 21st-century forest cover change. Science 342, 850–853. https://doi.org/10.1126/science.1244693.

Hansen/UMD/Google/USGS/NASA (2019). Global Forest Change 2000–2018. Data Download. Available on-line at http://earthenginepartners.appspot.com/science-2013-global-forest/download_v1.6.html.

Heinrich VH, R Dalagnol, HL Cassol, TM Rosan, CT de Almeida, CH Silva Junior, WA Campanharo, JI House, S Sitch, TC Hales, et al. (2021). Large carbon sink potential of secondary forests in the Brazilian Amazon to mitigate climate change. Nature communications 12, 1–11. https://doi.org/10.1038/s41467-021-22050-1.

Hijmans RJ (2019). raster: Geographic Data Analysis and Modeling. Available on-line at https://CRAN.R-project.org/package=raster.

Kalamandeen M, E Gloor, E Mitchard, D Quincey, G Ziv, D Spracklen, B Spracklen, M Adami, L. Aragão, and D Galbraith (2018). Pervasive rise of small-scale deforestation in Amazonia. Scientific reports 8, 1–10. https://doi.org/10.1038/s41598-018-19358-2.

Kuhn M, J Wing, S Weston, A Williams, C Keefer, A Engelhardt, T Cooper, Z Mayer, B Kenkel, the R Core Team, M Benesty, R Lescarbeau, A Ziem, L Scrucca, Y Tang, C Candan, and T Hunt. (2019). caret: Classification and Regression Training. R package version 6.0-84, available on-line at https://CRAN.R-project.org/package=caret.

LeSage J (2015). Spatial econometrics. In: Handbook of research methods and applications in economic geography. Edward Elgar Publishing. https://doi.org/10.4337/9780857932679.

Lloyd JD and YM León (2019). Forest change within and outside protected areas in the Dominican Republic, 2000-2016. bioRxiv. https://doi.org/10.1101/558346.

Mangiafico S (2019). rcompanion: Functions to Support Extension Education Program Evaluation. Available on-line at https://CRAN.R-project.org/package=rcompanion.

Myers R, J O’Brien, D Mehlman, and C Bergh (2004). Evaluación del manejo del fuego en los ecosistemas de tierras altas de la República Dominicana, Global Fire Initiative Informe Técnico. Tech. rep. The Nature Conservancy.

NASA (2019a). “MODIS Collection 6” standard quality Thermal Anomalies / Fire locations (MCD14ML), processed by the University of Maryland. Available on-line from the LANCE FIRMS operated by NASA’s Earth Science Data and Information System (ESDIS) platform at: https://earthdata.nasa.gov/firms. https://doi.org/10.5067/FIRMS/MODIS/MCD14DL.NRT.006.

NASA (2019b). VIIRS 375 m standard Active Fire and Thermal Anomalies product (VNP14IMGTML), processed by the University of Maryland. Available on-line from the LANCE FIRMS operated by NASA’s Earth Science Data and Information System (ESDIS) platform at: https://earthdata.nasa.gov/firms. https://doi.org/10.5067/FIRMS/VIIRS/VNP14IMGT_NRT.002.

OEA (1967). Reconocimiento y evaluación de los recursos naturales de la República Dominicana. Tech. rep. Washington, US: OEA, 1967.

Olson DM, E Dinerstein, ED Wikramanayake, ND Burgess, GV Powell, EC Underwood, J. D’amico, I Itoua, HE Strand, JC Morrison, et al. (2001). Terrestrial Ecoregions of the World: A New Map of Life on Earth: A new global map of terrestrial ecoregions provides an innovative tool for conserving biodiversity. BioScience 51, 933–938. https://doi.org/10.1641/0006-3568(2001)051[0933:TEOTWA]2.0.CO;2.

ONE (1982). 7mo. Censo Nacional Agropecuario 1982. Vols. 1 & 2. Tech. rep. Oficina Nacional de Estadística (ONE), Secretariado Técnico de la Presidencia.

ONE (2016). Precenso Nacional Agropecuario 2015. Informe de resultados. Tech. rep. Oficina Nacional de Estadística. Ministerio de Economía, Planificación y Desarrollo.

Ovalle de Morel E and A Rodríguez Liriano (1984). Análisis de la deforestación y la foresta en la República Dominicana. Eme Eme: Estudios Dominicanos. https://doi.org/ https://hdl.handle.net/20.500.12060/1341.

Pebesma E (2018). Simple Features for R: Standardized Support for Spatial Vector Data. The R Journal 10, 439–446. https://doi.org/10.32614/RJ-2018-009.

Pebesma E (2019). stars: Spatiotemporal Arrays, Raster and Vector Data Cubes. Available on-line at https://CRAN.R-project.org/package=stars.

QGIS Development Team (2020). QGIS. Available on-line at http://www.qgis.org/.

R Core Team (2020). R: A Language and Environment for Statistical Computing. Available on-line at https://www.R-project.org/. R Foundation for Statistical Computing. Vienna, Austria.

Sakamoto Y, M Ishiguro, and G Kitagawa (1986). Akaike information criterion statistics. Dordrecht, The Netherlands: D. Reidel 81. https://doi.org/10.1080/01621459.1988.10478680.

Sokal RR and NL Oden (1978). Spatial autocorrelation in biology: 1. Methodology. Biological journal of the Linnean Society 10, 199–228. https://doi.org/10.1111/j.1095-8312.1978.tb00013.x.

Tennekes M (2018). tmap: Thematic Maps in R. Journal of Statistical Software 84, 1–39. https://doi.org/10.18637/jss.v084.i06.

Tolentino L and M Peña (1998). Inventario de la vegetación y uso de la tierra en la República Dominicana. Moscosoa 10. Available on-line at https://www.biodiversitylibrary.org/item/181376, 179–203.

Tropek R, O Sedláček, J Beck, P Keil, Z Musilová, I Šímová, and D Storch (2014). Comment on “High-resolution global maps of 21st-century forest cover change”. Science 344, 981–981. https://doi.org/10.1126/science.1248753.

UN System Task Team on the Post-2015 UN Development Agenda (2012). Realizing the Future We Want for All. Report to the Secretary-General. United Nations New York, NY.

Venables WN and BD Ripley (2002). Modern Applied Statistics with S. Fourth. ISBN 0-387-95457-0. New York: Springer.

Wendell Werge R (1974). La agricultura de tumba y quema en la República Dominicana. Eme Eme: Estudios Dominicanos. https://doi.org/http://hdl.handle.net/20.500.12060/726.

Weston S (2019). foreach: Provides Foreach Looping Construct. Available on-line at https://CRAN.R-project.org/package=foreach.

Wickham H (2017). tidyverse: Easily Install and Load the ‘Tidyverse’. Available on-line at https://CRAN.R-project.org/package=tidyverse.

Zweifler MO, MA Gold, and RN Thomas (1994). Land Use Evolution in Hill Regions of the Dominican Republic. The Professional Geographer 46, 39–53. https://doi.org/10.1111/j.0033-0124.1994.00039.x.

